# ANKZF1 helps to eliminate stress-damaged mitochondria by LC3-mediated mitophagy

**DOI:** 10.1101/2024.11.05.622196

**Authors:** Mudassar Ali, Anjali, Koyeli Mapa

**Affiliations:** Protein Homeostasis Laboratory, Department of Life Sciences, School of Natural Sciences, Shiv Nadar Institution of Eminence, Delhi-NCR, Greater Noida, Gautam Buddha Nagar, Uttar Pradesh 201314, India

**Keywords:** Mitochondria, Mitophagy, proteotoxic stress, PINK, Parkin, LC3

## Abstract

Mitochondria, the double membrane-bound organelles of endosymbiotic origin, are crucial centers for cellular energy production and several essential metabolic pathways. Recent studies reveal that mitochondria become dysfunctional following numerous cellular stresses, and during pathologies, demanding an extensive investigation of mitochondrial turnover mechanisms. Apart from the specific response pathways to tackle different stresses, mitophagy or degradation of mitochondria by autophagy is a critical quality control mechanism that clears irreversibly damaged mitochondria. Mitophagy is majorly executed either by receptor-mediated or PINK1-Parkin-dependent pathways. Here, we show that the human orthologue of yeast Vms1, ANKZF1, participates in PINK-Parkin-mediated mitophagy. We show that ANKZF1 is extensively recruited to damaged mitochondria along with Parkin during mitochondrial proteotoxic stress induced by the expression of a single misfolded/aggregated protein or during uncoupler-induced membrane depolarization. Importantly, ANKZF1 recruitment to damaged mitochondria is significantly enhanced in the presence of Parkin, and ANKZF1 physically interacts with Parkin and LC3 during mitochondrial proteotoxic or depolarization stresses. ANKZF1 harbors six putative LC3-interacting regions (LIRs), LIR4 present at residues 333-336 is particularly important for ANKZF1-LC3 interaction. Furthermore, we show that *ANKZF1* knockout cells are compromised in clearing stress-damaged mitochondria by mitophagy, indicating an important role of ANKZF1 in mitochondrial turnover during stress. In summary, we show a new role of ANKZF1 in eliminating the stress-damaged mitochondria, reiterating the mito-protective role of Vms1/ANKZF1 during mitochondrial stresses.

## Introduction

Understanding organelle homeostasis under different cellular stresses has become one of the key areas of research in modern biology. Mitochondria is a major sub-cellular organelle that is frequently damaged and becomes dysfunctional during various cellular stresses and pathological conditions. Such stresses affect mitochondrial dynamics and functions, necessitating the elimination of damaged mitochondria and the subsequent biogenesis of new mitochondria to maintain the required pool of functional mitochondria in the cell.

Various small molecule stressors have been employed to induce mitochondrial stress to understand the response to such stresses. Mitochondrial membrane depolarizing agents like CCCP (Carbonyl Cyanide m-ChloroPhenylhydrazone) [1, 2], specific respiratory chain blockers e.g. rotenone [3, 4], oligomycin, antimycin, mitochondrial DNA damaging agents like ethidium bromide (EtBr) [5], or oxidizing agents that produce Reactive Oxygen Species (ROS) e.g. paraquat [1], H_2_O_2_ [4, 6, 7] have been extensively used as mitochondrial stressors. In some studies, specific mutant mitochondrial proteins were expressed within the organelle to understand the response pathways to mitochondrial protein misfolding or proteotoxic stresses. In metazoans like *C. elegans* and mammalian cells, the commencement of mitochondrial Unfolded Protein Response (mitoUPR) was reported upon DNA damage-induced stress [5] and proteotoxic stress, respectively. In some recent studies, protein import to mitochondria was blocked to induce stress in the organelle [8]. Many such studies revealed the existence of novel response pathways to manage stress in and around mitochondria. Recent works in yeast, *Saccharomyces cerevisiae,* revealed discoveries of pathways like Mitochondrial <u>C</u>ompromised <u>P</u>rotein import <u>R</u>esponse (mitoCPR) [9], Mitochondrial <u>T</u>ranslocation <u>A</u>ssociated <u>D</u>egradation (mitoTAD) [10], Unfolded Protein Response <u>a</u>ctivated by <u>m</u>istargeting of proteins (UPRam) [11–14], early mitoUPR [15], and Mitochondria Associated Degradation (MAD) [16, 17] etc. Recently, the contribution of the Ribosome Quality Control (RQC) pathway was shown to be vital for the protection of mitochondria from RQC byproducts like CAT (C-terminal Alanine Threonine extension)-tailed precursor proteins [18–20].

In case of overwhelming stress, damaged mitochondria are cleared by mitophagy. Upon activation, mitophagy is initiated by the dedicated receptor proteins forming phagophores. Mitophagy receptors or autophagy adaptor proteins initiate the phagophore formation by interacting with the Atg8-family of proteins like LC3 (Microtubule-associated protein 1A/1B-light chain 3) or GABARAPs (Gamma-Amino-Butyric Acid type 1 Receptor) present in the phagophore membrane that in turn initiate the autophagosome formation surrounding the damaged organelle. These phagophores ultimately encircle the damaged mitochondria, forming autophagosomes, and subsequently fuse to lysosomes forming autophago-lysosomes. Within the autophago-lysosomes, the lysosomal hydrolytic enzymes digest the engulfed material.

<u>P</u>TEN-<u>In</u>duced putative <u>K</u>inase protein 1 (PINK1) is involved in an alternate pathway of mitochondrial quality control and clearance [21–24]. Whenever PINK1 senses the mitochondrial membrane damage, it accumulates on the outer mitochondrial membrane where it further recruits Parkin, an E3 ubiquitin ligase from cytosol [1, 22, 25]. PINK1 phosphorylates ubiquitin and activates the E3 ubiquitin ligase activity of Parkin, which further governs the polyubiquitination of various outer mitochondrial membrane proteins on damaged mitochondria. These polyubiquitinated proteins are sensed by mitophagy adapter proteins, which act as a bridge between damaged mitochondria and autophagosomes. The adaptor proteins interact with polyubiquitinated proteins via their ubiquitin-associated (UBA) domain, on the other hand, they also interact with LC3 present on the phagophore membrane via the LIR (LC3 Interacting region) motif. These associations lead to the encircling of the damaged mitochondria by phagophores forming autophagosomes followed by clearance of the damaged organelle.

ANKZF1 (Ankyrin repeat and Zinc Finger Peptidyl tRNA Hydrolase 1) is the mammalian orthologue of Vms1 protein of yeast, *Saccharomyces cerevisiae.* Previous literature shows its involvement in the Ribosome Quality Control (RQC) pathway [26–29], where it acts as a tRNA hydrolase and recycles the nascent peptides from stalled ribosomes [26, 27, 29]. Importantly, Vms1 was shown to balance the CAT-tailing activity by RQC component Rqc2 and hence decreases the chance of import of CAT-tailed mitochondrial precursor proteins to the organelle reducing the chance of aggregation of CAT-tailed proteins within mitochondria [18–20]. Other studies showed the involvement of Vms1 in the recruitment of ubiquitin-proteasome system component Cdc48/VCP on damaged mitochondria during oxidative stress, to facilitate the Mitochondria-Associated-Degradation (MAD) pathway [16, 30, 31]. Interestingly, it was shown that during H_2_O_2_-induced oxidative stress, both Vms1 and ANKZF1 are localized to mitochondria although the necessity of mitochondrial localization of these proteins remains elusive [31, 32]. Recently, we have shown the specific modulatory role of *VMS1* during proteotoxic stress in the mitochondrial matrix in yeast. We showed that deletion of *VMS1* leads to aggravation of proteotoxicity in the mitochondrial matrix [33]. All these findings highlight the crucial roles of Vms1 during mitochondrial stress.

In the current study, we explored the role of the human protein ANKZF1 in mitochondrial stresses. We show that, during single misfolded-protein-induced mitochondrial proteotoxic stress or depolarization stress by CCCP treatment, ANKZF1 associates with mitochondria and co-localizes with Parkin. ANKZF1 also shows interaction with LC3 during mitochondrial stresses. Furthermore, sequence analysis of the protein revealed the presence of six putative LIR motifs; out of these six LIRs, LIR-4 harbored at amino acid residues 333-336 is indispensable for LC3-interactions. We further show that the residues 337-370 of ANKZF1 are indispensable for co-localization with ubiquitin and Parkin, indicating the presence of its ubiquitin-binding domain (UBA) in this segment. Lastly, knockout cells of *ANKZF1* exhibited compromised clearance of stress-damaged mitochondria by mitophagy which can be completely complemented by overexpression of wild-type ANKZF1. Importantly, we show that the t-RNA hydrolase deficient mutant of ANKZF1 (Q246L) [27], although deficient in its role in the Ribosome Quality Control pathway, interacts with Parkin and LC3 during stress like WT ANKZF1, indicating no overlap of its tRNA hydrolase with its role in mitophagy. Taken together, our study shows an important aspect of ANKZF1 in mitochondrial homeostasis; it plays an important role in the clearance of damaged mitochondria by LC3-mediated mitophagy during proteotoxic stress or depolarization stress of the organelle.

## Results

### ANKZF1 efficiently recruits to mitochondria during certain chemical-induced stresses in the organelle exclusively in the presence of Parkin

ANKZF1 is a multi-domain protein like its yeast orthologue, Vms1 (Figure 1A). From the N-terminus, the protein has a less defined N-terminal segment (NTS), followed by a Zinc-finger domain (ZnF), VLRF domain (Vms1-like Release Factor Domain) which also contains its putative MTD (Mitochondrial Targeting Domain). VLRF domain is followed by two Ankyrin repeats, a coiled-coil domain, and a VCP-interacting motif (VIM). To check any role of ANKZF1 in mitochondrial stresses, we expressed a C-terminally Green Fluorescent Protein (GFP) tagged version of the protein in HeLa cells and analyzed its cellular localization using imaging by confocal microscopy upon treatment with different mitochondrial stressors. In unstressed cells, ANKZF1-GFP does not show any distinct co-localization (Figure 1B) with the mitochondrial network. In the presence of various small molecule mitochondrial stressors like sodium Azide, H_2_O_2_, paraquat, rotenone, rapamycin and CCCP, there was no noticeable mitochondrial localization or recruitment of ANKZF1 on the mitochondrial network (Figure 1C and 1D).

**Figure 1.**
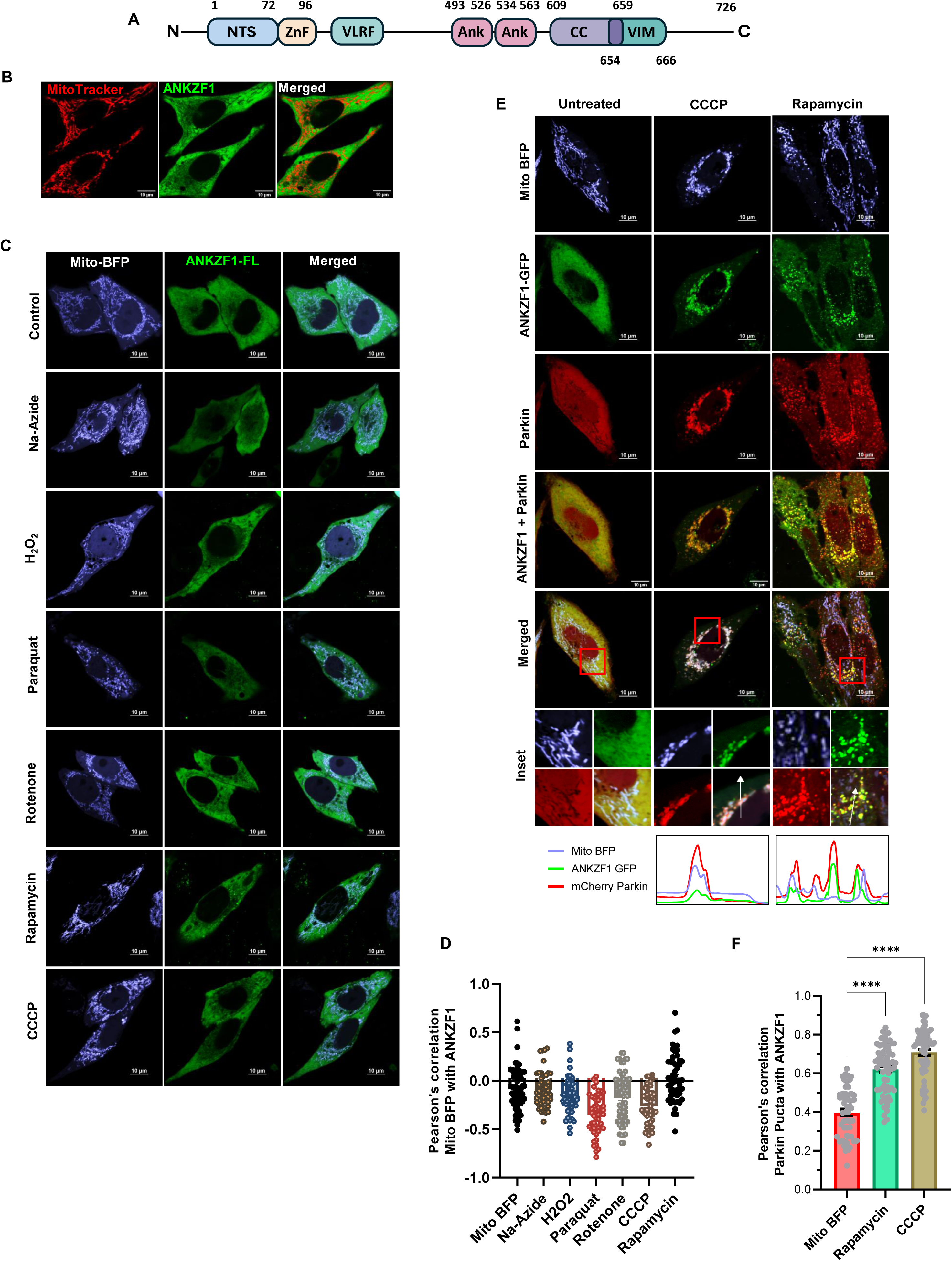
E3 ubiquitin ligase Parkin facilitates the mitochondrial recruitment of ANKZF1 on chemical stress-induced damaged mitochondria. **A.** Schematic representation of ANKZF1 domain map. From N-terminal segment of the protein has been shown as NTS, followed by a Zinc-finger domain (ZnF). After ZnF, there is VLRF (Vms1-Like Release Factor domain). VLRF is followed by two Ankyrin repeats (Ank), followed by a coiled-coil domain, a yet uncharacterized domain (shown as an orange box), and a VCP-interacting motif (VIM). The numbers mentioned indicate the number of amino acids in the primary sequence of the protein. **B.** Cellular distribution of ANKZF1-GFP is shown by imaging of transfected HeLa cells by confocal microscopy. MitoTracker-Red was used to stain the mitochondrial network. Right most panel is the merged panel of both red and green panels. **C.** ANKZF1 localization is shown during mitochondrial stress by various small molecule stressors (CCCP, H_2_O_2_, rotenone, Na-Azide, rapamycin and paraquat). In all stresses, ANKZF1 exhibits a diffused cytosolic localization except few punctate structures in rapamycin stress. **D.** Pearson’s correlation coefficient of ANKZF1-GFP co-localization with mito-BFP in the presence of chemical stressors. **E.** ANKZF1 is recruited as punctate forms on mitochondria during CCCP, and rapamycin-induced stress in the presence of Parkin overexpression in HeLa cells, Line-scan profile of fluorescence intensity of Mito-BFP, ANKZF1-GFP, and mCherry-Parkin are shown as blue, green, and red lines respectively, at the bottom of Rapamycin and CCCP treated condition. **F.** Pearson’s correlation coefficient of co-localization of mCherry-Parkin and ANKZF1-GFP was calculated in the non-stress control (Mito-BFP) conditions and during rapamycin and CCCP-treated conditions. Values represent means ± SEM, N = 3. Images of at least 50 cells were considered for quantification. Kruskal-Wallis test with Donn’s multiple comparison test was performed to determine the mean differences, **** indicates P < 0.0001.

CCCP treatment is known to cause mitochondrial stress and induce mitophagy majorly mediated by the PINK1-Parkin pathway [22, 34, 35]. As HeLa cells express Parkin at an insignificant level, we overexpressed Parkin with N-terminally-tagged mCherry and observed its association with mitochondria during chemical-induced mitochondrial stresses, as described above. As reported earlier, Parkin shows significant overlap with fragmented mitochondria post-CCCP treatment indicating recruitment of mitophagy apparatus on the depolarized and fragmented mitochondria (Figure 1E, middle column) [22]. Interestingly, when we co-expressed ANKZF1-GFP with Parkin, it showed significant co-localization with mCherry-Parkin indicating an interaction of ANKZF1 with Parkin (Figure 1E, middle column, and Figure 1F). This result reiterates the previously reported physical interaction between Parkin and ANKZF1 [36]. Similarly, upon rapamycin treatment, Parkin overlaps with the fragmented mitochondrial network, and ANKZF1 and Parkin are co-localized with fragmented mitochondria (Figure 1E, right column, and Figure 1F). With other stressors (rotenone, sodium-azide, H_2_O_2_ and paraquat), Parkin recruitment to the mitochondrial network remains insignificant, and ANKZF1 co-localization with mitochondria is not significant (Figure S1A).

To assess whether the mitochondrial membrane-depolarization is the key determinant of Parkin recruitment to the stressed mitochondria, we imaged the uptake of the mitochondrial membrane potential-dependent fluorescent dye, <u>T</u>etra<u>M</u>ethyl<u>R</u>hodamine <u>E</u>thyl ester (TMRE) [37], by mitochondria. Untreated mitochondria exhibited a bright TMRE signal completely overlapping with the mitochondrial network indicating intact mitochondrial membrane potential, as expected (Figure S1B, topmost row). CCCP treatment significantly diminished the TMRE uptake due to membrane depolarization (Figure S1B, second row from top and S1C). Other stressors like rapamycin, rotenone, paraquat and sodium-azide also showed significantly decreased uptake of TMRE, albeit less prominent than CCCP treatment, indicating defects in the maintenance of membrane polarization upon treatment with these compounds (Figure S1B and S1C). Notably, treatment with H_2_O_2_ showed normal TMRE uptake like untreated healthy cells (Figure S1B and S1C). Although most of these stressors led to membrane depolarization, at least partially, and mitochondrial fragmentation, Parkin and ANKZF1 recruitment to mitochondria were not significant. This result hints that the interaction between Parkin/ANKZF1 and mitochondria is stress-specific.

### Proteotoxic stress due to expression of mitochondria-targeted misfolded proteins leads to severe mitochondrial fragmentation and may lead to depolarization

We have recently shown that yeast Vms1 plays an important modulatory role during mitochondrial matrix proteotoxic stress [33]. To investigate the role of the mammalian orthologue of Vms1, ANKZF1, during mitochondrial proteotoxic stress, we developed a mitochondria-specific proteotoxic stress model in human cells. To impart proteotoxic stress exclusively to mitochondria, we targeted two exogenous model misfolded proteins: i) Parkinson’s disease-associated A53T mutant of α-synuclein (referred to as A53T-Syn hereafter) [38, 39] and ii) recently described amyloid-forming aggregation-prone protein, PMD (<u>P</u>rotein with <u>M</u>isfolded <u>D</u>omains) [33], to mitochondria. Specific targeting of model proteins was achieved using the <u>M</u>itochondrial <u>T</u>argeting <u>S</u>ignal (MTS) of SMAC (<u>S</u>econd <u>M</u>itochondria-derived <u>A</u>ctivator of <u>C</u>aspase) (Figure 2A) and the proteins were expressed as fusion proteins with a C-terminal GFP tag (Figure 2A). Confocal microscopy of the GFP-tagged proteins reveals complete overlap with the signal of the Mito-mCherry, indicating correct localization of the stressor proteins to mitochondria (Figure 2B and 2C). Mitochondria-targeted GFP, without any fused stressor proteins, serves as the control of overexpression of exogenous proteins within mitochondria. As reported before, PMD protein forms prominent amyloids in vitro [33]. When expressed within mitochondria, the protein forms SDS-insoluble aggregates and it is difficult to detect the expression of the protein by western blot using the standard method of denaturation for sample loading (Figure S2A). With exclusive use of lysis buffer containing a strong denaturant like 8M urea, PMD aggregates are solubilized, and the protein is detectable by western blot at the expected size, indicating formation of the SDS-insoluble aggregates by the protein in mitochondria (Figure S2B). A53T-Syn is detected by standard western blot techniques at the expected size (Figure S2A). Expression of both the stressor proteins leads to increased fragmentation of the mitochondria as evidenced by a significant reduction in the mitochondrial branch length (Figure 2B, 2D, 2E, and S2C), more prominently in the case of PMD protein-induced proteotoxic stress (Figure 2B, 2D and 2E).

**Figure 2.**
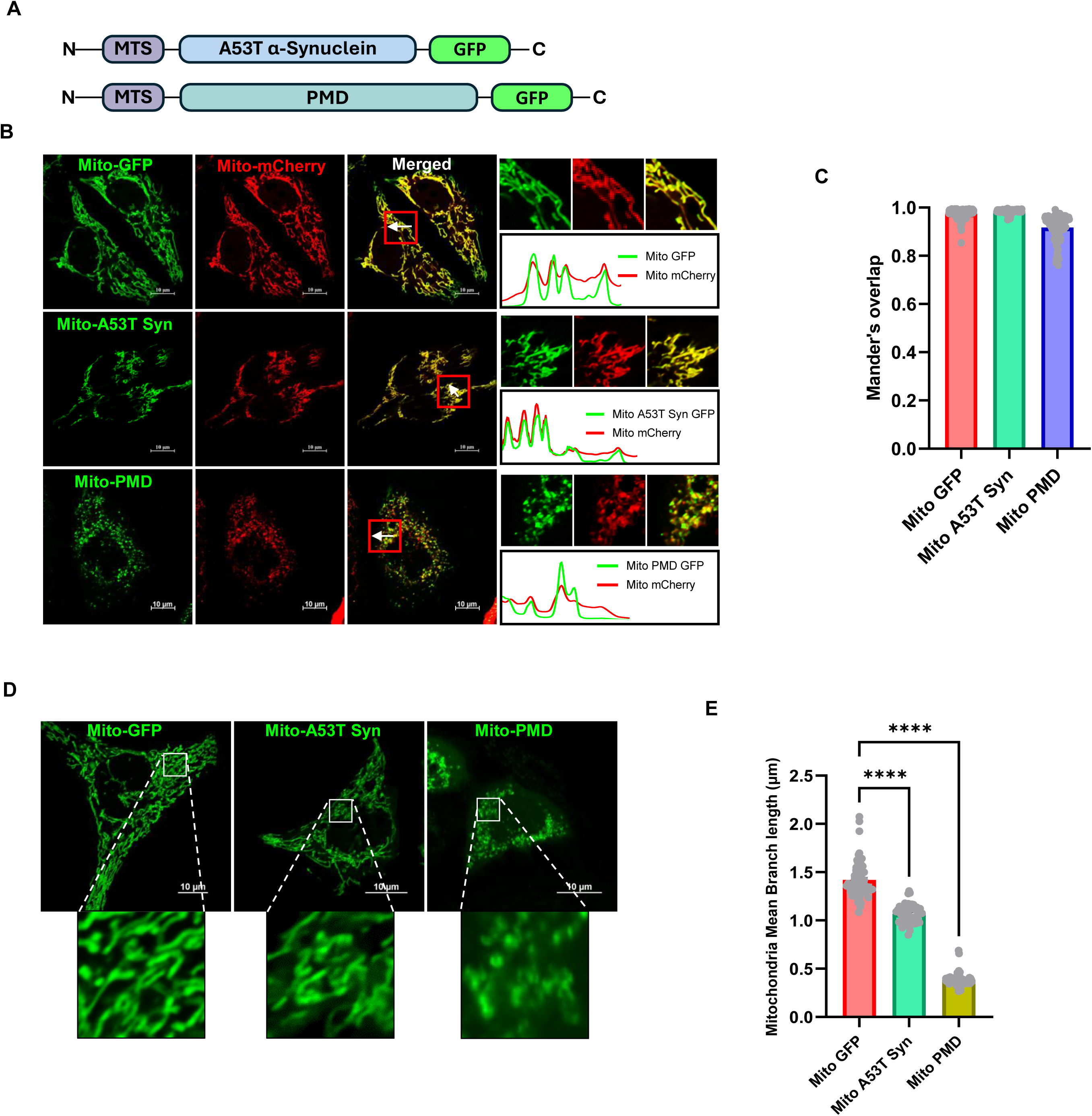
Generation of model of mitochondria-specific single misfolded/aggregated protein-induced proteotoxic stress, in human cells. **A.** Schematic representation of model stressor proteins, A53T α-synuclein, and PMD (Protein with Misfolded Domain)-fused to GFP, targeted to mitochondria. MTS denotes Mitochondrial Targeting Sequence. **B.** Expression and localization of mitochondria-targeted A53T α-synuclein and PMD proteins in HeLa cells were checked by imaging using confocal microscopy. Mitochondria-targeted mCherry (Mito-mCherry) was used as a mitochondrial marker Line-scan profile of fluorescence intensity of mitochondria-targeted GFP-tagged proteins (Mito-GFP, Mito-A53T Syn-GFP, and Mito-PMD-GFP) and Mito-mCherry are shown in green and red lines, respectively. **C**. Co-localization analysis of GFP-tagged control/stressor protein with Mito-mCherry by Mander’s overlap shows more than 90% co-localization score of these stressor proteins to mitochondrial marker, exhibiting correct mitochondrial targeting. Values represent means ± SEM (N = 3). **D and E.** Expression of both of these stressor proteins in mitochondria leads to increased mitochondrial fragmentation. However, PMD expression leads to significantly enhanced mitochondrial fragmentation, where the size of mitochondria is reduced to less than one-third of the length of healthy mitochondria. Values in panel E represent mitochondria mean branch lengths in µm ± SEM (N=3). As data did not follow a normal distribution, a non-parametric Kruskal Wallis test with Donn’s multiple comparison test was performed to determine the mean differences in both plots C and E, ****P < 0.0001.

Next, we investigated the mitochondrial membrane potential after imparting the proteotoxic stress, Mitochondrial TMRE uptake remained mostly unaffected in mitochondria expressing A53T-Syn mutant protein indicating intact membrane polarization despite stress due to the expression of this protein (Figure S3A and S3B). In contrast, expression of PMD protein in mitochondria led to severe depolarization (like CCCP treatment as shown in Figure S3A, bottom row) as indicated by significant defects in the uptake of TMRE compared to the control cells (Figure S3A and S3B). This data indicated that the extent of stress and its effect on the mitochondrial forms and function depends on the property of the misfolded protein.

### ANKZF1 interacts with mitophagy/autophagy machinery during proteotoxic stress in mitochondria

As proteotoxic stress leads to severe mitochondrial fragmentation as well as depolarization with PMD-induced stress, we went ahead to check whether the fragmented and damaged mitochondria are cleared by the autophagy pathway. First, we took the tandemly fluorescently tagged-LC3 (tfLC3) tagged with monomeric RFP (Red Fluorescent Protein) followed by GFP (Green Fluorescent Protein) [40] (Figure 3A) as a reporter protein. tfLC3 indicates flux of ongoing autophagy by differential pH sensitivity of GFP and monomeric RFP in the acidic milieu of the lysosome, in the final step of autophagy [40]. The GFP fluorescence of the tfLC3 is quenched in the strong acidic pH of lysosome after the fusion of the autophagosomes (containing GFP-RFP-LC3 in its membranes) to lysosomes. In contrast, the fluorescence of the monomeric RFP remains intact in the same acidic environment. Thus, during autophagy, when the autophagosomes fuse with lysosomes, only the red fluorescence of the autophagic cargo remains intact, and hence, the number of red puncta clearly outnumbers the green puncta. When we performed imaging of the tfLC3 using confocal microscopy in the control non-stressed cells, very few LC3 puncta were visible with both green and red fluorescence which completely overlapped with each other upon merging the images from both the channels [same number of puncta in the red channel and merged (yellow) channel] (Figure 3A, top row). When we co-expressed the tfLC3 with the mitochondrial stressor proteins (either A53T-Syn or PMD), the number of LC3 puncta increased (Figure 3A, middle and lower rows) compared to the control condition (Figure 3A, top row). Notably, the number of LC3 puncta is prominently more in PMD-protein-induced stress compared to stress due A53T-Syn expression, again indicating differential stress response depending on the nature of the protein that misfolds or aggregates within mitochondria. Interestingly, apart from the increased number of LC3 puncta, the number of puncta exclusively with RFP fluorescence was significantly high during proteotoxic stress (Figure 3A, middle and lower rows and Figure 3B). This data indicated that proteotoxic stress in mitochondria leads to increased bulk autophagy.

**Figure 3.**
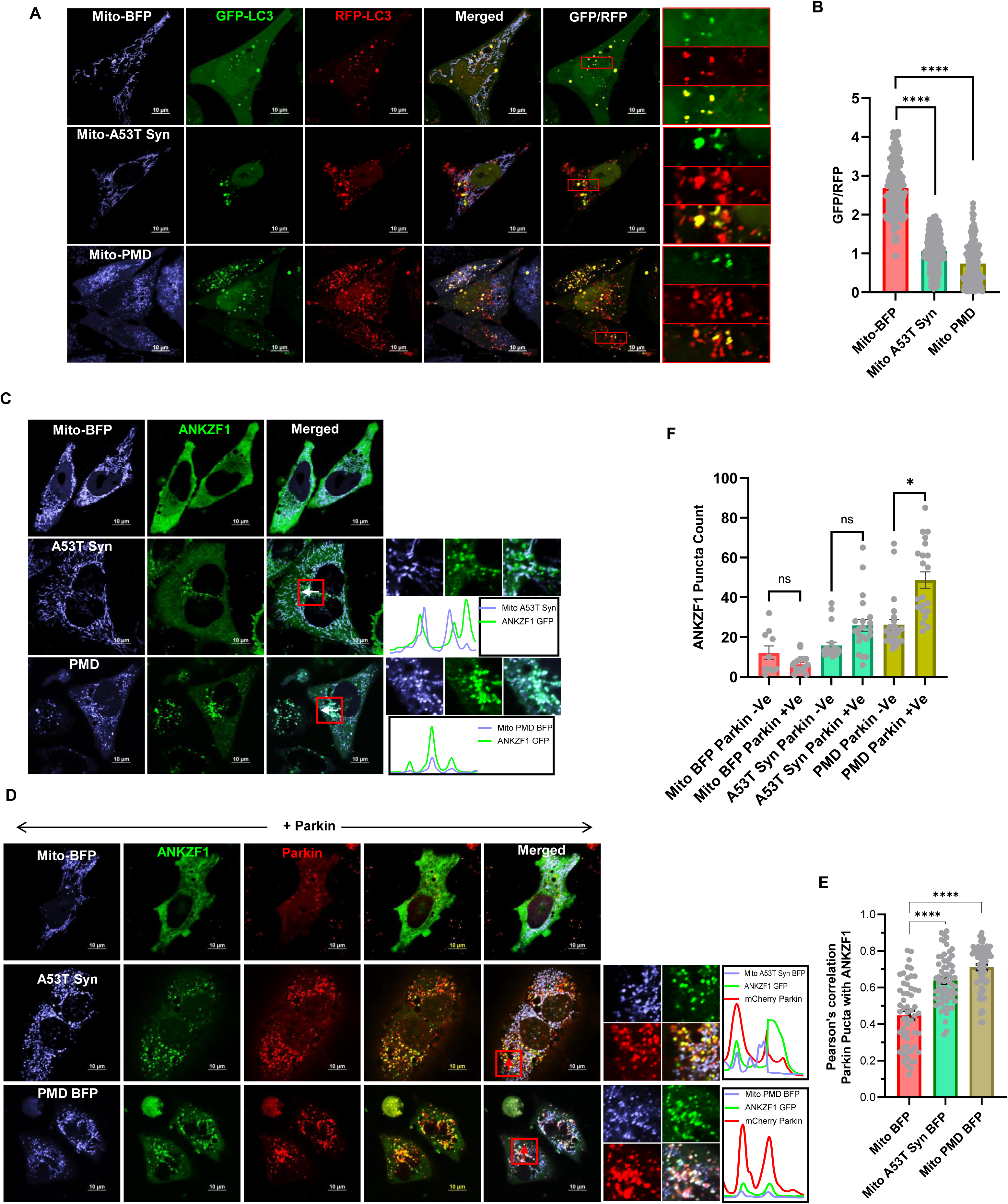
Parkin also assists in the recruitment of ANKZF1 on severely fragmented mitochondria during mitochondrial proteotoxic stress. **A-B.** Any changes in autophagic flux due to proteotoxic stress in mitochondria by expressing the stressor proteins, A53T α-synuclein or PMD, was further confirmed by expressing the fluorescent reporter protein mRFP-GFP tandem fluorescently-tagged LC3 (tfLC3). GFP and RFP fluorescence intensity of LC3 puncta were measured and GFP/RFP of each puncta were calculated and plotted as bar plots (B), as explained in the main text, mitochondrial proteotoxic stress leads to reduced GFP/RFP ratio of tfLC3 suggesting significantly increased overall autophagic flux during mitochondrial targeted proteotoxic stress in comparison to the control condition. Values represent means ± SEM (N= 3). As data did not follow a normal distribution, non-parametric Kruskal Wallis test with Donn’s multiple comparison test was performed to determine the mean differences, ****P < 0.0001**. C and D.** ANKZF1-GFP was co-expressed with A53T α-synuclein-BFP or PMD-BFP in the absence (panel C) or in the presence of mCherry-Parkin (panel D) in HeLa cells. Without any stress (control protein BFP is expressed in mitochondria), ANKZF1 shows diffused cytosolic localization (Upper row panel C). ANKZF1 forms some punctate foci during A53T-α-synuclein (middle row panel C) or PMD-induced stress (lower row panel C) in mitochondria. Line-scan profile of fluorescence intensity of BFP-tagged stressor proteins (Mito-A53T Syn BFP and Mito-PMD BFP) and ANKZF1-GFP are shown in blue and green colors, respectively (panel C). Similarly, in the presence of Parkin, along with blue and green lines, mCherry-Parkin fluorescence is represented as a red line (panel D) showing merged peaks suggesting co-localization. ANKZF1 puncta are recruited over mitochondria during mitochondrial proteotoxic stress even in the absence of Parkin (panel C). However, the recruitment of ANKZF1 is significantly increased when Parkin is overexpressed in HeLa cells (Panel D), suggesting ANKZF1 recruitment on stressed mitochondria is facilitated by Parkin. **E.** Pearson’s correlation coefficient of co-localization was calculated and plotted for Parkin and ANKZF1 puncta during unstressed control (Mito-BFP) condition, Mito-A53T Syn-BFP and Mito-PMD-BFP induced stress conditions, Values represent means ± SEM, (N= 3), data were not following normal distribution, Kruskal Wallis a non-parametric test with Donn’s multiple comparison test was performed to determine the mean differences, ****P < 0.0001. **F.** ANKZF1 puncta count per cell in the absence and presence of Parkin during control conditions (mitochondria expressing Mito-BFP) and proteotoxic stress conditions (mitochondria expressing Mito-A53T Syn-BFP and Mito-PMD-BFP), shows a significant increase in ANKZF1 puncta formation in presence of Parkin during proteotoxic stress condition. Values represent means ± SEM, (N= 3), data were not following normal distribution, Kruskal Wallis test with Donn’s multiple comparison test was performed to determine the mean differences, ns is non-significant, *P < 0.05.,

As described earlier, Parkin is one of the key regulators of PINK1/Parkin-mediated mitophagy, we monitored Parkin puncta recruitment to mitochondria to understand the extent of clearance of the damaged mitochondria during proteotoxic stress in the organelle by mitophagy. Upon expressing the stressor proteins, A53T-Syn or PMD, Parkin showed punctate structures mostly co-localized with the mitochondrial network (Figure S3C, middle and bottom rows and S3D), however in unstressed cells, Parkin exhibits diffused cytosolic localization (Figure S3C, top row, and S3D). The extent of Parkin puncta co-localized to the mitochondrial network was more prominently visible during PMD-induced mitochondrial proteotoxic stress (Figure S3C, bottom row, and S3D).

Upon checking the ANKZF1 cellular distribution (in the absence of Parkin) during mitochondrial proteotoxic stress, we found some punctate localization of ANKZF1 during stress (Figure 3C, middle and bottom rows) in contrast to diffused cytosolic localization of the protein in the control cells (Figure 3C, upper row). This was in contrast to chemical-induced stresses where the protein does not recruit to mitochondria at all, in the absence of Parkin. Upon overexpression of Parkin, ANKZF1 recruitment on mitochondria during proteotoxic stress was significantly increased (Figures 3D, 3E and 3F). As previously shown in the case of CCCP treatment, during proteotoxic stress also, ANKZF1 and Parkin puncta showed significant co-localization (Figures 3D and 3E). As ANKZF1 showed strong interaction with Parkin in multiple mitochondrial stresses, it indicated a possible role of the protein in the process of mitophagy.

To check the association of ANKZF1 with other players of mitophagy, its interaction with other components was further checked. During proteotoxic stress in mitochondria, ANKZF1 puncta showed significant co-localization with LC3 puncta over fragmented mitochondria (Figures 4A and 4B) reiterating its association with mitophagy/autophagy components. Co-expression of ANKZF1, Parkin and LC3 in the cells experiencing PMD-induced stress in mitochondria, shows complete co-localization of ANKZF1 with Parkin and LC3 in a ring-like structure (Figure 4C, lower panels). Interestingly, all three proteins are co-localized to each other during mitochondrial proteotoxic stress. This association was further checked for endogenous ANKZF1 protein, cells were co-transfected with Mito-PMD-BFP, GFP-Parkin and RFP-LC3 followed by ANKZF1 immuno-staining with anti-ANKZF1 antibody. In this case too, images show similar patterns of co-localized events of Parkin, LC3 and ANKZF1 during proteotoxic stress (Figure S4A). Furthermore, to determine the physical interaction of ANKZF1 with Parkin and LC3 during mitochondrial proteotoxic stress (by expressing PMD) and depolarization, we performed co-immunoprecipitation experiments. HEK293T cells were transfected with ANKZF1-GFP and mCherry-Parkin having mitochondrial stress due to expression of PMD or due to CCCP treatment. Mito-BFP was used as a control of exogenous folded protein overexpression in mitochondria. Importantly, Parkin was exclusively co-precipitated with anti-GFP antibody, indicating a physical interaction between ANKZF1-GFP and Parkin during PMD-induced proteotoxic stress as well as mitochondrial depolarization stress due to CCCP treatment (Figure 4D). In the control condition (with Mito-BFP expression), no interaction between Parkin and ANKZF1 was observed. Endogenous LC3 was precipitated with ANKZF1-GFP during PMD-induced proteotoxic stress and during CCCP-induced stress (Figure 4D), indicating an interaction between LC3 and ANKZF1, as observed by imaging experiments. Furthermore, ANKZF1’s interaction with lysosome was also checked during proteotoxic stress. ANKZF1 showed a significant co-localization with lysosomal protein LAMP1, suggesting fusion of ANKZF1-containing autophagosomes with lysosomes indicating the role of the protein in mitophagy (Figure S4B and S4C).

**Figure 4.**
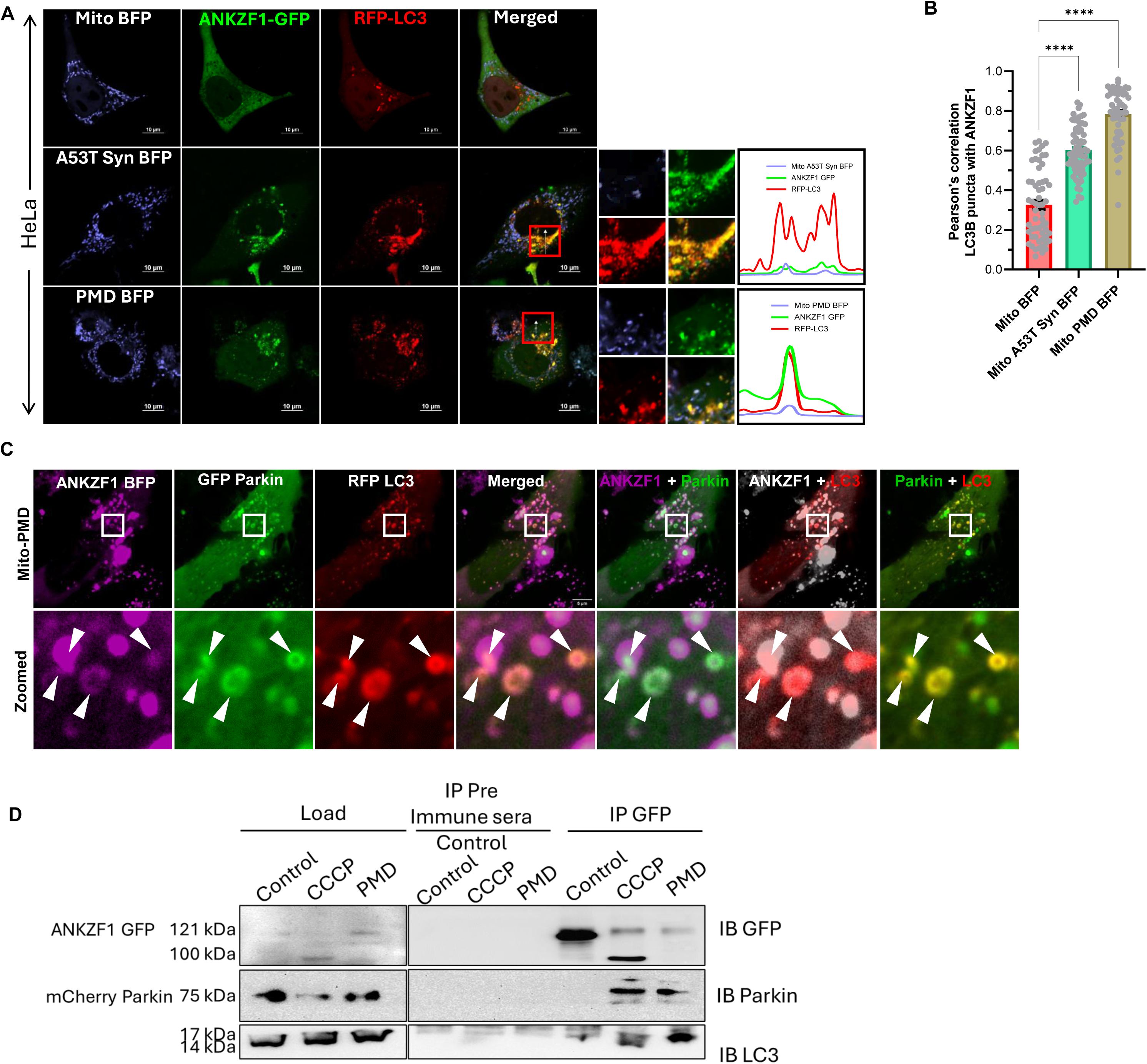
ANKZF1 shows interaction with Parkin and LC3 during mitochondrial proteotoxic stress and CCCP-induced stress. **A.** ANKZF1-GFP and RFP-LC3 were co-expressed in HeLa cells along with Mito-BFP, A53T-α-synuclein-BFP, and PMD-BFP. In the control panel (Mito-BFP), ANKZF1 shows diffused cytosolic expression, and low LC3 puncta formation was observed, but in stressed conditions, lower two panels (A53T α-synuclein BFP and PMD BFP) a high number of RFP-LC3 puncta formation and ANKZF1 puncta formation was observed. Line-scan profile of fluorescence intensity of BFP-tagged protein constructs (Mito-A53T Syn BFP, and Mito-PMD BFP) targeted to mitochondria shown in blue and ANKZF1 GFP shown in green and RFP-LC3 was indicated in the red color profile. **B.** Pearson’s correlation coefficient of co-localization was calculated for RFP-tagged LC3 puncta and GFP-tagged ANKZF1 during unstressed control (Mito-BFP), Mito-A53T-Syn-BFP, and Mito-PMD-BFP conditions. Values represent means ± SEM, N=3, data were not following normal distribution, Kruskal Wallis test (a non-parametric test) with Donn’s multiple comparison test was performed to determine the mean differences, ****P < 0.0001. **C.** ANKZF1-BFP, GFP-Parkin, and RFP-LC3 were co-expressed along with Mito-PMD, showing puncta of ANKZF1, Parkin, and LC3 are getting co-localized. The zoomed panel shows the highly co-localized phagophore like structures of ANKZF1, Parkin, and LC3. **D.** Co-immunoprecipitation of mCherry-Parkin and LC3 with ANKZF1-GFP was performed during CCCP-treated depolarization and PMD-induced proteotoxic stress conditions. Left panel shows the 5% load of whole cell lysate before binding to Protein A/G Sepharose beads, while right panel shows immunoprecipitated proteins (IP) with pre-immune antisera and anti-GFP antibody. Blots were probed with anti-GFP (top), anti-Parkin (middle) and anti-LC3 (bottom) antibodies (Supplementary Table 2).

### ANKZF1 possesses features of mitophagy adaptor proteins

ANKZF1’s interaction with Parkin and LC3 during mitochondrial stresses (proteotoxic and depolarization) indicated its possible role as a mitophagy adaptor protein. As reported before, all mitophagy receptor or autophagy adaptor proteins harbor a particular amino acid sequence consisting of aromatic amino acids (W/Y/F) followed by any two amino acids (XX) and end with an aliphatic amino acid (L/I/V) [41]. These regions of the autophagy/mitophagy adaptor proteins are known as <u>L</u>C3-Interacting <u>R</u>egion (LIR) [42–45]. When we checked the human mitophagy receptor/adaptor proteins, NIX, BNIP3, p62, optineurin, or AMBRA1, we found the conserved LIR motif in all the known mitophagy receptors/adaptors, as expected (Figure 5A). Interestingly, we found six such putative LIR motifs in the human ANKZF1 protein sequence (Figure 5B). After multiple sequence analyses of ANKZ1 sequences of different mammals, we found most of them contain multiple such conserved LIR sequences (Figure 5C). To check the importance of these predicated LIRs for ANKZF1-LC3 interaction, we first made three truncation mutants of ANKZF1; Δ210-ANKZF1 (containing deletion of N-terminal 210 amino acids), Δ330-ANKZF1 (containing deletion of N-terminal 330 amino acids) and Δ370-ANKZF1 (containing deletion of N-terminal 370 amino acids) (Figure 5D). As shown in Figure 5D, the Δ210 mutant is devoid of LIR-1 and LIR-2, Δ330 mutant is deleted of LIR-1, LIR-2, and LIR-3. Δ370 mutant lacks the first 5 LIRs (LIR-1 to LIR-5) and contains only the LIR-6 (495^th^-498^th^ residues). Interestingly, Δ210-ANKZF1 and Δ330-ANKZF1 truncation mutants retain the co-localization with LC3 puncta, like wild-type ANKZF1 protein during PMD-induced stress in the mitochondria (Figures 5E, 5F), suggesting that LIR-1, LIR-2 and LIR-3 are dispensable for interaction with ANKZF1. In contrast, Δ370-ANKZF1 did not show any punctate structure co-localized with LC3 during the same stress, indicating an important role of LIR-4, LIR-5 and LIR-6 in LC3 interaction (Figures 5E, 5F). This experiment was recapitulated in SHSY-5Y cells and similar findings were observed (Figure S5A and S5B)

**Figure 5.**
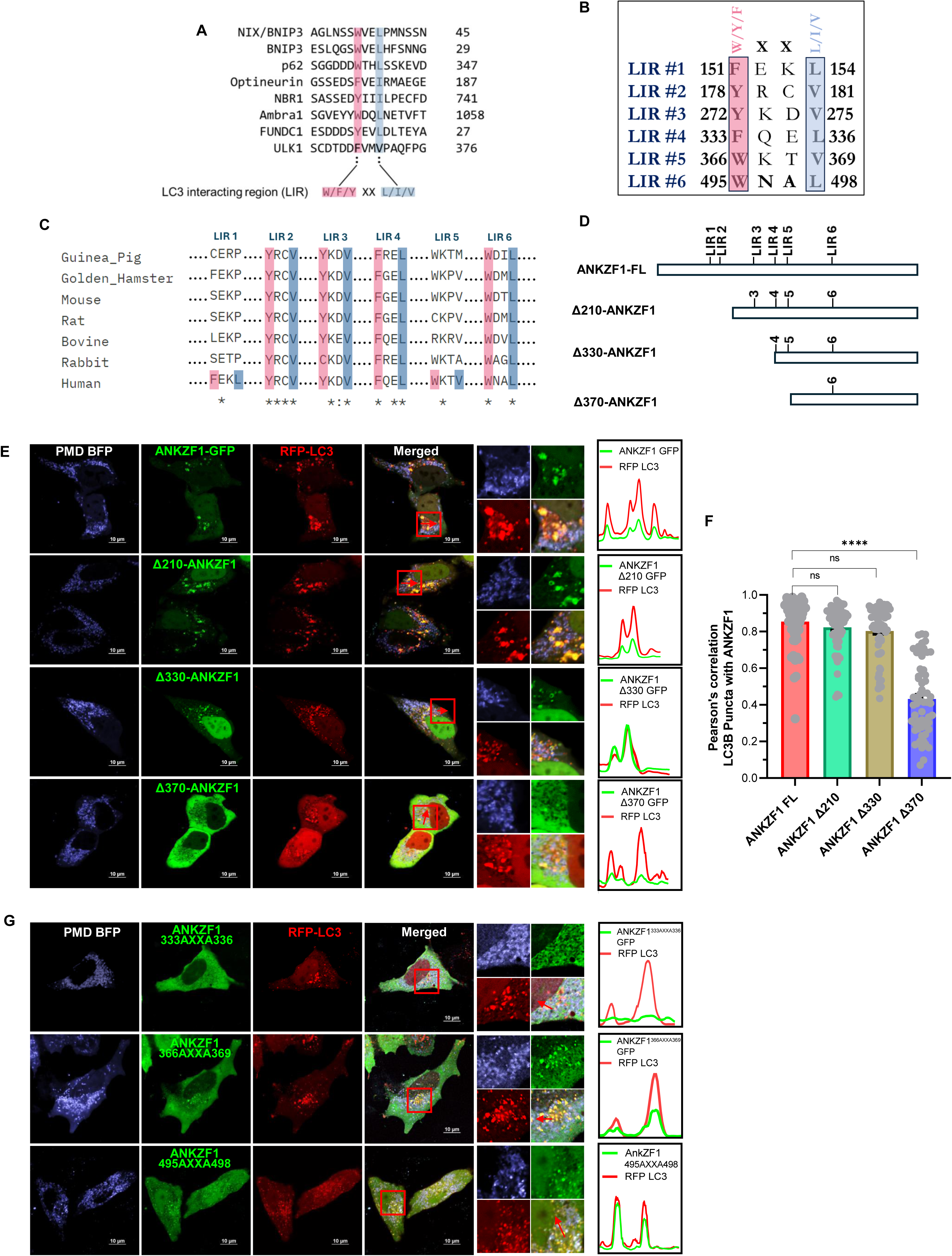
ANKZF1 sequence contains six putative LC3-Interacting Regions (LIRs), and LIR-4 (333-336 residues) is indispensable for LC3-interaction. **A.** Multiple sequence analysis (MSA) of some of the known mitophagy/autophagy adaptor/receptor proteins shows the presence of a conserved LIR (LC3 interacting region) motifs. B. ANKZF1 sequence analysis shows the presence of six different probable LIR motifs consisting of similar sequence like known adaptors/receptors, named as LIR1 to LIR6. **C.** MSA of ANKZF1 sequence from different mammals shows the presence of conservation of LIR2, LIR3, LIR4 and LIR6. **D.** Schematic overview to depict the distribution of 6 LIR-like motifs in ANKZF1 full-length protein sequence, three truncation mutants of ANKZF1 (Δ210, Δ330, Δ370) were designed to screen the probable functional LIR motifs crucial for interaction with LC3. Schematic representation of truncation mutants show the presence and absence of LIRs in these three mutants compared to the full-length wild type ANKZF1 protein (ANKZF1 FL). **E.** ANKZF1 recruitment and interaction with LC3 during PMD-induced mitochondrial proteotoxic stress were checked after expressing the truncation mutants of ANKZF1. ANKZF1-FL shows punctate foci that co-localize with RFP-LC3 puncta during proteotoxic stress (panel E, upper row). Δ210-ANKZF1, Δ330-ANKZF1 truncation mutants also show similar behavior and significantly co-localize with LC3 puncta (panel E, second and third rows from the top) like the full-length protein. Δ370-ANKZF1 shows a diffused cytosolic pattern indicating no interaction with LC3. Line-scan profile of fluorescence intensity of RFP-LC3 puncta and WT-ANKZF1/ Δ210-ANKZF1/Δ330-ANKZF1 puncta shows merged intensity peak suggesting co-localization. In contrast, Δ370-ANKZF1-GFP puncta and RFP-LC3 puncta peaks do not merge, suggesting the absence of interaction. ANKZF1 peaks are shown in green and LC3 peaks are shown in red. **F.** Pearson’s correlation coefficient of co-localization was calculated and plotted for LC3 puncta and ANKZF1 (WT-ANKZF1, Δ210-ANKZF1, Δ330-ANKZF1, and Δ370-ANKZF1) puncta during Mito-PMD BFP induced stress conditions, Values represent means ± SEM, N=3, data were not following normal distribution, Kruskal Wallis a non-parametric test with Donn’s multiple comparison tests was performed to determine the mean differences, ****P < 0.0001. **G**. ANKZF1 LIR mutants, F333A-L336A, W366A-V369A, and W495A-L498A fused with GFP were co-transfected with RFP-LC3 in HeLa cells along with PMD-BFP. W366A-V369A and W495A-L498A mutants of ANKZF1 show interaction and co-localization with RFP-LC3 like the wild type ANKZF1 (middle and lower panel), F333A-L336A mutant of ANKZF1 during PMD induced stress is not showing any co-localization with RFP-LC3 (top panel). Line-scan profile of fluorescence intensity of mutant ANKZF1-GFP and RFP LC3 puncta shows merged intensity peak suggesting co-localization.

Furthermore, to confirm the exact LIR required for LC3 interaction, we generated point mutations by site-directed mutagenesis in the LIR-4, LIR-5 and LIR-6 (Figure S7C). We mutated the W/Y/F and L/I/V residues to Alanine. Thereafter, the co-localization of mutant ANKZF1 with LC3 was analyzed during PMD-induced proteotoxic stress. Notably, mutating the LIR-5 and LIR-6 of ANKZF1 did not abolish its interaction with LC3. However, LIR-4 (amino acids 333-336) mutant showed no co-localized events between ANKZF1 and LC3 (Figure 5G). This result indicates the importance of LIR-4 (333 FQEL 336) in the interaction with LC3. Thus, LIR-4 is the critical LIR motif in the ANKZF1 required for interaction with LC3 to facilitate mitophagy.

We further checked the interaction of LIR-4 mutant of ANKZF1 with Parkin during proteotoxic stress and found that both proteins are co-localized (Figure S5C and S5D). Thus, LIR-4 is specifically required for interaction with LC3.

As ANKZF1 shows prominent co-localization with Parkin on stressed mitochondria and ANKZF1’s recruitment to stress-damaged mitochondria is significantly enhanced in the presence of Parkin, it is likely that ANKZF1 should interact with ubiquitinated mitochondrial proteins. Mitophagy adaptors are of varied nature structurally and so far, both mitochondrial constituent proteins as well as some cytosolic proteins, have been shown to act as mitophagy adaptors [46, 47]. The presence of a ubiquitin-associated domain (UBA) is reported for most of the cytosolic mitophagy adaptor proteins as well as some mitochondrial integral proteins that act as adaptors. Thus, next, we asked whether ANKZF1 contains a ubiquitin-binding domain. We checked the ANKZF1 interaction with ubiquitin during PMD-induced stress and found ANKZF1 puncta co-localizes with Ub-DsRed puncta, suggesting that it possesses a ubiquitin-binding motif (Figure S5E and S5F). To further delineate the position of the putative ubiquitin-binding-domain of ANKZF1, we checked ANKZF1 puncta formation in the presence of Parkin (Figure S6A and S6B) as well as ANKZF1-puncta that co-localize with Ub-DsRed (Figure S6C and S6D) using a panel of truncation mutants of ANKZF1. As there are no well-conserved motifs reported for UBA-like domains for mitophagy adaptors, we checked a number of N-terminal truncation mutants for this purpose. As ubiquitin-binding domains are usually located following the LIRs in the primary sequences of the adaptors, we took Δ330-ANKZF1, Δ370-ANKZF1, Δ410-ANKZF1 and Δ450-ANKZF1 truncation mutants and compared the puncta formation capacity with respect to the WT protein. Interestingly, only Δ330-ANKZF1 showed punctate co-localization with Parkin (Figure S6A and S6B) and Ub-DsRed (Figure S6C and S6D) during PMD-induced proteotoxic stress similar to WT ANKZF1. Deletion of N-terminal 370 amino acids or more residues, abolished the ANKZF1-puncta formation and co-localization with Parkin and Ub-DsRed. It is important to reiterate that ANKZF1 showed puncta formation co-localized to Ub-DsRed even in the absence of Parkin overexpression, albeit to a lower extent (Figure S6C and S6D). From this data, we can conclude that the ubiquitin-binding domain of ANKZF1 is located after residue 331 downstream of LIR and residues 337 to 370 are indispensable for ubiquitin-binding. As we were not confirmed about the exact location of the ubiquitin-binding domain, we checked the predicted structure of the 337-450 residues. Intriguingly, this part of the protein is predicted to form α-helical structures by I-TASSER [48, 49] (Figure S7A), as reported for UBA-domains of the mitophagy adaptors. Thus, we have designated residues 337-450 as the putative UBA domain of ANKZF1 (Figure S7B).

In conclusion, we found that residues 333-336 ANKZF1 constitute the functional LIR of the protein, and for the ubiquitin-binding, 337-370 residues are indispensable. These residues predominantly form an α-helical domain as reported for other UBA-domains of mitophagy adaptors. Thus, ANKZF1 possesses all major criteria for mitophagy adaptor proteins and helps in clearance of damaged mitochondria by LC3-mediated mitophagy.

### Generation and characterization *ANKZF1* knockout (KO) cell lines

To check any impairment in autophagic clearance of damaged mitochondria in the absence of ANKZF1, we generated the CRISPR-Cas9 mediated knockout of *ANKZF1* in HeLa cells (Figure 6A). We used two separate pairs of sgRNAs for the deletion of *ANKZF1* (Figure 6A). *ANKZF1* KO was confirmed by PCR (Figure S8B) and Sanger sequencing (Figure S8A) of the edited genomic locus, which shows the deletion of ∼3kb fragment from the gene sequence (Figure S8A and S8B). The deletion was further validated by western blot, confirming no production of the ANKZF1 protein in the KO lines (Figure 6B). For all experiments, two clones of *ANKZF1* KO cells (clone 1 and clone 2) were used (Figure 6A, 6B, Figure S8A, S8B). After generating the *ANKZF1* KO cells, we checked for any changes that the KO cells may encounter in the absence of ANKZF1, especially in terms of mitochondrial health. We checked TMRE-based mitochondrial membrane potential and found that *ANKZF1* deletion does not lead to any significant changes in the mitochondrial membrane potential (Figures 6C and 6D). Next, we assessed mitochondrial morphology by measuring the mitochondrial branch length. Mitochondrial branch length remained unchanged in *ANKZF1* KO cells in the absence of any stress (Figures 6E, top row, and 6F). Even after imparting proteotoxic stress, the reduction in mitochondrial branch length in *ANKZF1* KO cells was similar to the WT HeLa cells (Figure 6E, middle and bottom rows, and Figure 6F). We further checked PINK1 and Parkin recruitment status on the mitochondria in *ANKZF1* KO cells during no-stress control conditions (Figure S8C, left panel) and during proteotoxic stress (Figure S8C, right panel). In the control condition, no aberrant recruitment of PINK1 or Parkin on mitochondria was observed in the *ANKZF1* KO cells, indicating no significant stress in mitochondria and lack of induction of mitophagy exclusively due to the absence of ANKZF1 (Figure S8C). Although *ANKZF1* KO cells did not exhibit any significant mitochondrial stress, the viability of both clones of KO cells was significantly decreased upon cycloheximide (CHX) treatment compared to WT cells (Figure S9A). This result further validates the KO of ANKZF1 by showing missing functions of ANKZF1 in the Ribosome Quality Control (RQC) pathway during translation block by CHX treatment. Furthermore, we performed complementation by checking the restoration of cell viability by overexpressing WT ANKZF1 during CHX treatment and compared it with a previously described t-RNA hydrolase-deficient mutant (RQC-deficient) of ANKZF1 (Q246L) [27]. Importantly, cell viability could be fully restored by expressing the WT ANKZF1 protein but not by the Q246L mutant of ANKZF1 during CHX treatment (Figure S9A).

**Figure 6.**
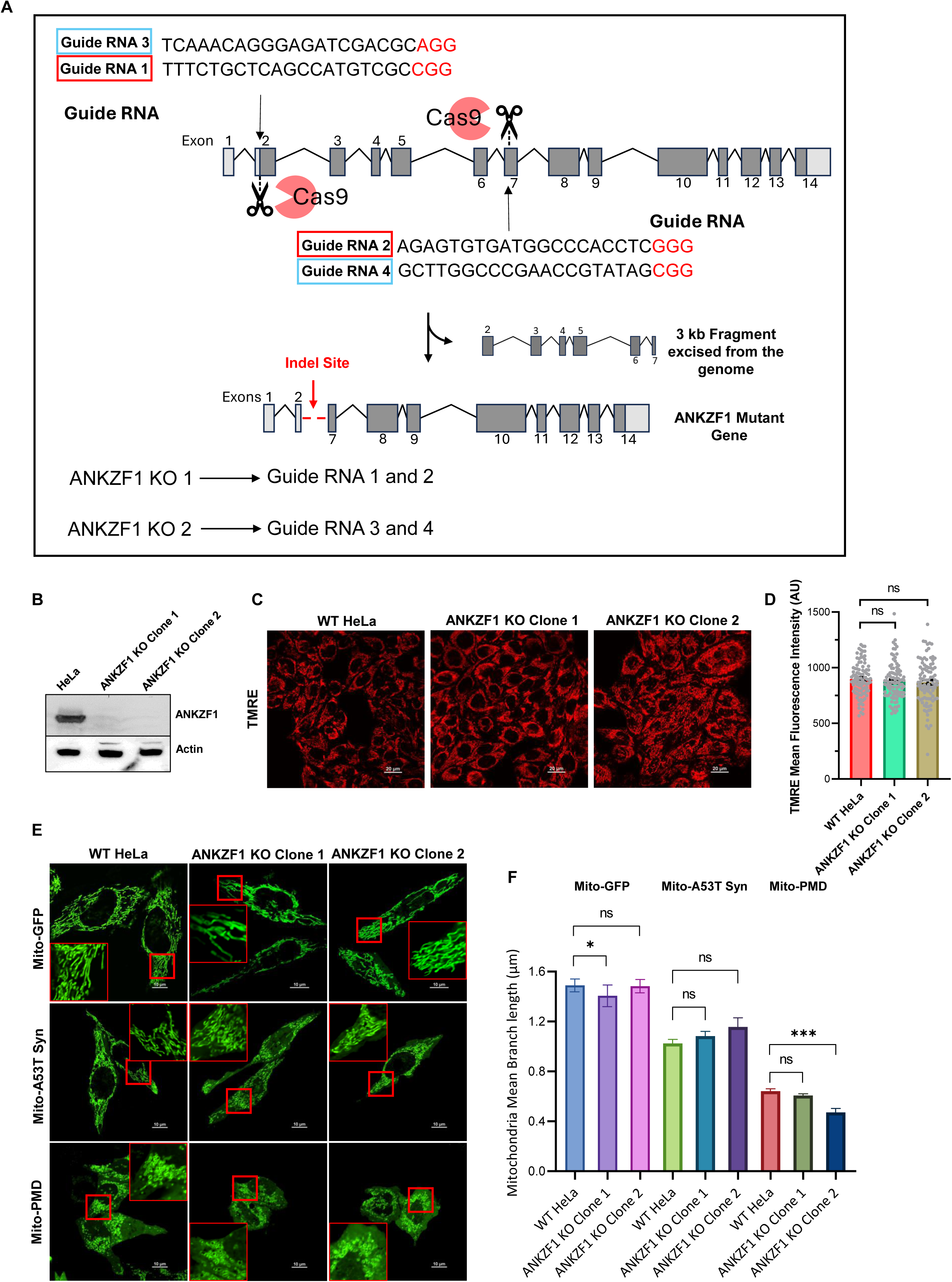
ANKZF1 knockout (KO) cell generation and its characterization. **A.** Schematic representation of CRISPR-Cas9 mediated *ANKZF1* knockout (KO) generation. The sequence of two sets of guide RNAs (sgRNAs) that were used in combination is shown in the schematic. **B.** Confirmation of *ANKZF1* knockout cells by western blot with ANKZF1-specific antibody. Actin was used as a loading control. **C.** Wild-type HeLa cells and *ANKZF1* KO clones were stained with TMRE to assess the mitochondrial membrane potential. **D.** TMRE mean fluorescence intensity plot of wild-type HeLa and *ANKZF1* KO cells. Values represent means ± SEM, N=3. Data were not following normal distribution, Kruskal Wallis a non-parametric test with Donn’s multiple comparison test was performed to determine the mean differences. **E.** Mitochondrial morphology in *ANKZF1* KO cells was compared with wild-type HeLa cells in healthy control cells and in the presence of proteotoxic stressor proteins A53T-Synuclein-GFP and PMD-GFP expressed in mitochondria. **F.** Mitochondrial mean branch length was measured in wild-type HeLa and both the *ANKZF1* KO clones. Graphs show overall no significant changes in the mitochondrial mean-branch length due to deletion of *ANKZF1* gene. Values represent means ± SEM, N=3, as data were not following normal distribution, Kruskal Wallis (a non-parametric test) with Donn’s multiple comparison test was performed to determine the mean differences, *P < 0.05, ***P < 0.001.

To check whether a compromised function of ANKZF1 in the RQC pathway also leads to any change in its role in mitophagy, we checked the recruitment of the ANKZF1-Q246L mutant during proteotoxic stress in mitochondria. We found ANKZF1-Q246L mutant is similarly recruited to stress-damaged mitochondria like the WT protein in the presence of Parkin overexpression. ANKZF1-Q246L also exhibited significant co-localization with Parkin (Figures S9B and S9C) and LC3 (Figures S9D and S9E) in both the KO clones as well as in the WT cells, suggesting that t-RNA hydrolase activity of ANKZF1 does affect its interaction with Parkin and LC3.

### ANKZF1 knockout (KO) cells are specifically defective in mitophagy, but bulk cellular autophagy remains unaffected

To assess any alterations in mitophagy in the absence of ANKZF1, we utilized the Mito-Keima (Mt-Keima) reporter described before [50–52], in the *ANKZF1* KO cells. Mt-Keima was co-expressed along with control protein or stressor protein PMD in mitochondria of WT HeLa cells and *ANKZF1* KO cells to evaluate any alterations in the mitophagy events in the absence of ANKZF1. The images of Mt-Keima captured at 488 nm excitation and 640 nm emission are depicted in green and the images taken after excitation at 560 nm and emission at 640 nm are shown in red (Figure 7A). In WT HeLa cells, as expected, the average mitophagy events (red signal to green signal ratio) during mitochondrial proteotoxic stress (due to expression of PMD) were found to be extremely high (Figure 7A and S10A, 2nd row from top) in comparison to the non-stressed control cells (Figure 7A and S10A, uppermost row). Although *ANKZF1* KO cells also showed higher mitophagy events during PMD-induced mitochondrial stress (Figures 7A and S10A, third and fourth rows from top), when we compared the shift of Keima488 to Keima560 (change of mitochondrial pH from basic/neutral to acidic), we found significant difference between WT and KO cells, suggesting reduced mitophagy in KO cells (Figure 7B and S10B). However, this change was reverted to WT level when ANKZF1 was overexpressed in the KO cells (Figure S10B and S10C). Also, the average number of mitophagy events between WT and KO cells expressing PMD, we observed a significant reduction in the average mitophagy events in the KO cells (Figure 7C). We further checked if cells lacking *ANKZF1* are impaired in bulk autophagy or not, we used the tf-LC3 reporter as described in the earlier sections. For that purpose, tf-LC3 was expressed with either mito-BFP (control) or mito-PMD protein in WT and KO cells. Increased autophagy was observed after PMD expression compared to control cells (Figure S10D). However, no significant difference in bulk autophagy was observed between WT and *ANKZF1* KO cells (Figure S10D) suggesting ANKZF1 is specifically important for mitophagy but not for bulk autophagy.

**Figure 7.**
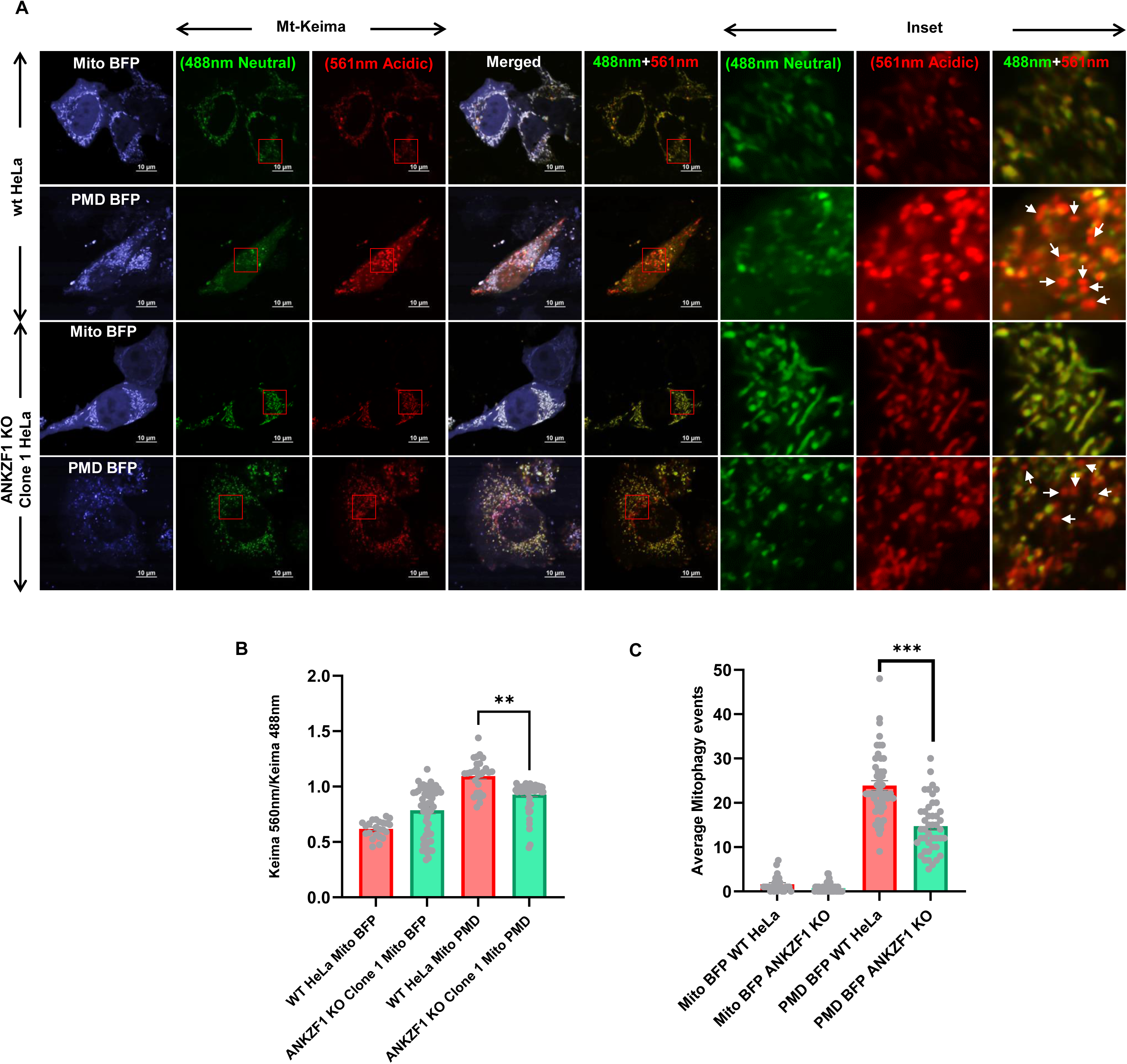
Cells deleted of *ANKZF1* show compromised mitophagy during mitochondrial proteotoxic stress. **A.** The mitophagy efficiency of WT HeLa cells and *ANKZF1* KO cells were compared by using pH dependent Mt-Keima protein as explained in the results. KO cells showed functional mitophagy events but significantly decreased mitophagy events in comparison to WT HeLa cells during PMD-induced mitochondrial proteotoxic stress. **B.** Ratiometric analysis of Keima560 vs Keima488 fluorescence intensity, PMD expressed WT-Hela cells showing high 560/488 ratio in comparison to the KO cells corresponding high mitophagy rate. **C.** Average mitophagy events per cell during control and stressed conditions in WT-HeLa cells and *ANKZF1* knockout cells were assessed by counting the red signals (autophago-lysosome) from the merged panel of Mt-Keima (panel A). Red-only vesicles are shown with white arrowheads in the inset of the merged panel (rightmost column of panel A). Values represent means ± SEM, (N =3). As data were not following a normal distribution, Kruskal Wallis a non-parametric test with Donn’s multiple comparison test was performed to determine the mean differences, **P < 0.01, ***P < 0.001.

Next, alterations in mitophagy events were also checked in *ANKZF1* KO cells by inducing mitochondrial depolarization by CCCP treatment. The ratio of Mt-Keima fluorescence emission after excitation at 560nm with the emission after excitation at 488 nm was used as a surrogate for ongoing mitophagy as described before. In WT cells, CCCP treatment led to an increase in the ratio due to increased mitophagy (Figure S11A, top two rows). In the *ANKZF1* KO cells, Mt-Keima ratio was significantly reduced compared to WT cells following CCCP treatment indicating ineffective mitophagy in the absence of ANKZF1 (Figures S11A, and S11B).

In summary, we show that the absence of ANKZF1 leads to a prominent reduction in clearance of stress-damaged mitochondria by mitophagy although the bulk cellular autophagy remains unaltered in the absence of protein. Thus, ANKZF1 plays a crucial role in LC3-mediated mitophagy to clear out irreversibly damaged mitochondria.

## Discussion

Mitophagy or specific clearance of damaged or surplus mitochondria by the process of autophagy, is a complex multi-step process involving various components. There are several mechanisms by which the process of mitophagy can take place; in receptor or adaptor-mediated mitophagy where the mitophagy adaptor proteins, usually some outer mitochondrial proteins, interact with LC3 of growing phagophore membrane and initiate the engulfment of mitochondria within autophagosomes [46, 47]. In the PINK1-Parkin-dependent pathway, classically, the depolarized mitochondria accumulate non-translocated PINK1 on the mitochondrial surface, which helps in the recruitment of Parkin, an E3 ubiquitin ligase on the mitochondrial outer membrane. Parkin helps in ubiquitination of outer mitochondrial membrane proteins, which interact with the mitophagy adaptor/receptor proteins. These adaptor and receptor proteins act as bridges between the mitochondrial outer membrane and LC3 on the growing phagophore membrane, thus facilitating the encircling of the damaged mitochondria by isolation membranes, ultimately forming autophagosomes. Thus, mitophagy adaptor proteins are crucial players in the whole course of autophagic clearance of mitochondria from the early steps of the process. So far, many mitophagy adaptor proteins have been discovered, and some of these proteins have been identified only recently [53, 54].

In this study, we show that human ANKZF1 (yeast orthologue of Vms1), a known component of the mammalian Ribosome Quality Control (RQC) pathway, helps in the removal of stress-damaged mitochondria by mitophagy. Our data reveals that ANKZF1 prominently recruits to damaged mitochondria in the presence of Parkin and interacts with LC3 through its LIR and co-localizes with ubiquitin indicating interaction with ubiquitin during mitochondrial stress and thus fulfils all the criteria of a mitophagy adaptor protein.

### Varied nature of mitophagy adaptor proteins

Among the known mitophagy adaptor proteins, many outer mitochondrial membrane proteins and some cytosolic proteins are known to get recruited to mitochondrial surface to initiate the process of mitophagy, when the need arises [46, 47]. Recently, inner mitochondrial proteins like Prohibitin2 [54] and matrix proteins like Bag6 [53] were also shown to act as mitophagy adaptor proteins. Mitochondria-resident proteins that work as mitophagy adaptors can be involved in receptor-mediated mitophagy without the involvement of PINK1-Parkin-like machinery, as these adaptors can directly interact with LC3 with their LIRs and initiate the process of mitophagy. For inner membrane proteins like Prohibitin, the protein localizes to the mitochondrial outer membrane upon damage of the organelle and outer membrane disruption [54]. Thus, adaptor proteins involved in receptor-mediated mitophagy do not necessarily require the presence of any ubiquitin-interacting domains or motifs. Recently discovered mitophagy adaptors like Prohibitin or Bag6 are not reported to possess a UBA-like domain [53, 54]. For PINK-Parkin-dependent mitophagy, the adaptors usually rely on interacting with ubiquitinated mitochondrial proteins and such adaptors usually contain a ubiquitin-binding domain. Well-established mitophagy adaptor/receptor proteins like p62, Optineurin, NDP52, NBR1, and AMBRA1 interact with the autophagosomes via a conserved LIR motif, on the other end, these adaptors also interact with polyubiquitinated outer mitochondrial membrane proteins through specific ubiquitin-binding domains (UBD). UBDs typically form an alpha-helical structure to facilitate interaction with the β-sheet of ubiquitin. However, the ubiquitin-binding domains of these adaptor proteins do not share well-defined conserved motifs or sequences, suggesting the diverse nature of the ubiquitin-binding domains. Apart from the α-helix forming UBD, there are Zinc finger-associated UBD (UBZ) or Pleckstrin homology domain (PH) known to interact with polyubiquitinated proteins [55]. The type of UBDs may vary from protein to protein; for example, p62 and NBR1 contain a UBA, Optineurin contains a UBAN domain, and AMBRA1 contains a WD40 domain. Thus, inherently the ubiquitin-binding domains are structurally more diverse in contrast to LIRs which contain a conserved motif.

In this work, we delineated the LIR motif of ANKZF1 which is indispensable for interaction with LC3. We also demonstrated the interaction of ANKZF1 with ubiquitin during mitochondrial stress and narrowed down the segment of protein essential for interaction with ubiquitin. Although the segment of the protein essential for ubiquitin-binding is predicted to be α-helical, the exact position and nature of the ubiquitin-binding domain remain to be elucidated.

We have summarized the role of ANKZF1 as a mitophagy adaptor protein in Figure 8 during mitochondrial stress. We have shown the various steps how stress-damaged mitochondria can be cleared by ANKZF1 by LC3-mediated mitophagy (Figure 8). This finding reiterates the importance of this protein in the protection of mitochondria in various ways, either acting in the mito-RQC pathway or as a mitophagy adaptor protein.

**Figure 8.**
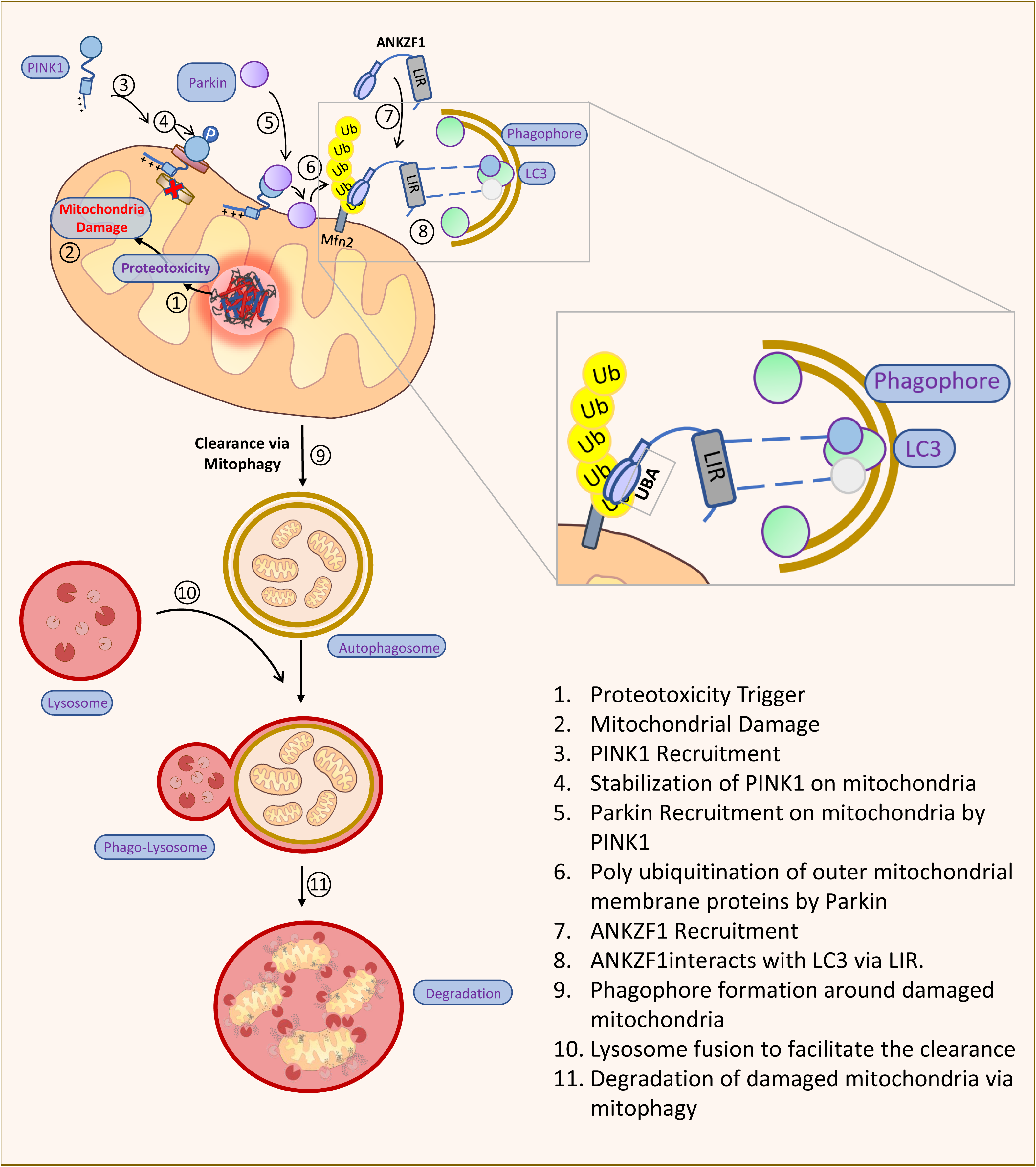
An illustrated summary of ANKZF1’s role as a mitophagy adaptor protein. Figure 8 shows a summary of ANKZF1’s role as mitophagy adaptor protein. The box shows the interaction of LC3-Interacting Regions (LIR) of ANKZF1 with LC3 on the growing phagophore membrane. The flow diagram on the left shows the stepwise (steps 1-11) process of mitophagy induction and clearance of damaged mitochondria during proteotoxic stress in the organelle. Here, ANKZF1 has been shown to act as a mitophagy adaptor, which helps in autophagosome formation surrounding the damaged mitochondria by interacting with ubiquitinated mitochondrial outer membrane proteins through its putative UBA domain and LC3 on the growing phagophore membrane.

### ANKZF1, a known component of the mammalian RQC pathway, also plays a crucial role in LC3-mediated mitophagy during specific mitochondrial stresses

Here, we showed that ANKZF1 recruits to mitochondria during some specific stresses like mitochondrial depolarization by CCCP or rapamycin-treatment. We also checked the role of ANKZF1 during mitochondrial stress by single aggregation-prone misfolded proteins as previous literature reported that many proteins, especially neurodegenerative-disease-associated aggregation-prone cytosolic proteins like α-synuclein [56], mutant huntingtin [57] or amyloid β-peptide [58] that are found in mitochondrial sub-compartments and lead to mitochondrial stress and dysfunction. Our model of mitochondria-specific proteotoxic stress model mimics such stress arising from protein misfolding and aggregation. Importantly, by using two different model aggregation-prone proteins as stressor protein we show that the extent of mitochondrial fragmentation and loss of membrane potential are dependent on the protein being misfolded or aggregated within the organelle. The proteotoxic stress generated by specific aggregated proteins like PMD, is sufficient to activate the clearance of damaged mitochondria by mitophagy. It is noteworthy to mention that some misfolded and aggregation-prone proteins (like A53T-Syn) do not cause membrane depolarization, yet the stress imparted to mitochondria by these proteins is sufficient to send signals for the onset of mitophagy.

Furthermore, using our stress model, we could segregate ANKZF’1’s role in RQC from its role in mitophagy. We showed that the tRNA-hydrolysis deficient mutant (RQC mutant) of ANKZF1 similarly interacts with Parkin and LC3 like the WT protein during mitochondrial stress indicating the intact function of the protein in mitophagy, despite loss of tRNA hydrolase function.

The major question remains; is there any advantage in engaging an RQC component in mitophagy? Recent literature shows the importance of eIF5A, a translation initiation factor that also plays a multifactorial role in RQC. eIF5A prevents ribosome-pausing on polyproline-containing proteins like Tim50, which is crucial for mitochondrial biogenesis and function [59]. Besides, eIF5A also plays an important role in CAT-tailing [60]. Very recent literature shows that eIF5A is the main component behind mitochondria-associated protection of ubiquitinated aggregation-prone proteins from proteasomal degradation [61]. From all this evidence, it is tempting to speculate that ANKZF1, being a component of RQC, would be in close proximity to eIF5A, which shelters the mitochondria-associated ubiquitinated aggregation-prone proteins from degradation. These aggregated and ubiquitinated proteins on mitochondria could be sensed by ANKZF1. In case of an overwhelming accumulation of mitochondria-associated aggregation-prone proteins, a preferred mechanism to restore the proteostasis would be removal of the organelle with associated aggregation-prone proteins by initiating mitophagy over degradation of aggregated proteins from the mitochondrial surface. A mitophagy adaptor, which is also a component of RQC, would be a reliable candidate for sensing the mitochondrial stress for and initiation of mitophagy in such a scenario. However, the triggering factor for ANKZF1 recruitment to damaged mitochondria remains to be explored in the future.

## Material and Methods

### Cell Culture, Transfection and Treatments

HeLa and HEK293T cells were cultured in DMEM (low glucose) containing 10% FBS (Gibco) and maintained at 37°C with 5% CO_2_. SHSY-5Y cells were cultured in 45% DMEM, 45% Ham’s F-10 media supplemented with 10% FBS, and maintained at 37°C with 5% CO_2_. All transfection of plasmid DNA in HeLa and SHSY-5Y cells was done by lipofectamine 2000^TM^ (Invitrogen) and polyethylenimine (PEI) was used for plasmid transfection in HEK293T cells using the manufacturer’s recommended protocols. Cells were administered with 30µM, 60µM, and 90µM of the uncoupler Carbonyl Cyanide m-ChloroPhenyl hydrazone (CCCP) to disrupt the mitochondrial membrane potential. To induce bulk autophagy, cells were treated with 200nM Rapamycin for 24 hours. To increase the cellular ROS production, cells were treated with 2mM Paraquat or 1mM H_2_O_2_ for 12 hours and 2 hours respectively. To induce mitochondrial damage, cells were treated with 5µM rotenone for 4 hours and 5mM Sodium Azide for 3 hours. To stain the mitochondria, cells were treated with 200nM MitoTracker deep red for 30 minutes before imaging.

### Plasmids and Cloning

Constructs used in this study include the following: EBFP2-N1 (Addgene, # 54595), mCherry Parkin (Addgene), mCherry-Mito-7 (Addgene #55102), ptfLC3 (Addgene #21074), Mito GFP, RFP-LC3, pHAGE-mt-mKeima (Addgene #131626). ANKZF1-FL-GFP, Δ210-ANKZF1-GFP, Δ330-ANKZF1-GFP, and Δ370-ANKZF1-GFP constructs were generated by using standard restriction digestion-based molecular cloning technique, and desired genes were ligated with digested pEGFPN1 (Clontech) empty vector backbone. ANKZF1 LIR mutants F333A-L336A, W366A-V369A, and W495A-L498A were generated by overlap-PCR based site-directed mutagenesis and cloned into pEGFPN1 vector. Mito-PMD-GFP construct was previously made in our laboratory and the cloning method was explained previously [33]. Mito-A53T-α-Synuclein-GFP constructs were generated in the lab by using the same methodology where A53T α-synuclein gene was amplified from EGFP-alphasynuclein-A53T (Addgene #40823) and N-terminally fused with mitochondria targeting sequence of SMAC (Second Mitochondrial derived Activator of Caspase) by overlap PCR followed by the cloning of the overlapped fragment in digested pEGFPN1 vector. Mito-PMD-BFP and Mito-A53T-α-Synuclein-BFP constructs were generated by sub-cloning into EBFP2-N1 (Addgene #54595). LAMP1 DsRed construct was generated by subcloning of LAMP1 from LAMP1-emiRFP670 (Addgene #136570) into mCherryN1 plasmid backbone, plasmids were digested with XhoI and EcoRI restriction enzymes, followed by the ligation and transformation. Similarly, UbG76VdsRed construct was generated by subcloning of UbG76V from UbG76VGFP (Addgene # 11941) into DsRedN1 vector backbone, both the vectors were digested with EcoRI and BamHI restriction enzymes, followed by the ligation and transformation by standard protocol. Q246L ANKZF1 mutant was generated in the lab by PCR-based side-directed mutagenesis method in the p-EBFP-N1 and p-EGFP-N1 vector backbone.

### Live Cell Imaging and Image Analysis

All microscopy images were acquired using an Apochromat 100X 1.4 NA oil immersion-based objective lens in a Nikon A1R MP+ Ti-E confocal microscope system. Imaging was performed at temperature (37°C), CO_2_ (5%) and humidity controlled conditions. 405nm, 488nm, and 560nm lasers were used to excite the signals from BFP, GFP, and RFP/mCherry channels respectively, and the emission signals were detected by an automated du4 detector. mt-mKeima imaging was done by using the manual detector, where 488nm and 560nm lasers were used to excite the signals of neutral and acidic environments respectively, while both the emissions were collected at 620nm. Clearly visible ANKZF1 puncta and mt-mKeima puncta were manually counted in FIJI (NIH). TMRE and tf-LC3 fluorescence intensity and co-localization assessment were analysed and calculated in Nikon NIS Essentials analysis software.

Mitochondria mean branch length was measured by using ImageJ plugin MiNa (mitochondrial network analysis). Initially, raw images were opened in ImageJ and then following processing pipeline was used to process the raw image before analysis. ***Step 1****. Process – Noise - Despeckle, **Step 2**, Process – Enhance – CLACHE, **Step 3**, Plugins – Analyze – Tubeness, **Step 4**, Process – Binary – Make Binary, **Step 5,** Process – Binary – Skeletonize, **Step 6a**, Plugin – Mitochondria Analyzer – 2D – 2D Analysis, **Step 6b**, Plugins – Stuart Lab – MiNa Script – MiNa Analyzer Morphology.* Steps 6a and 6b both give a similar range of mitochondrial branch length, which was further plotted.

### Immunofluorescence Assay

Cells were seeded on the coverslip 24 hours before the transfection. Mito PMD-BFP, GFP-Parkin and RFP-LC3 were transfected, and cells were incubated for 24 hours. Next day, media were discarded, and cells were washed twice with 1XPBS and fixed with 4% formaldehyde at room temperature for 20 minutes, followed by permeabilization and blocking by 0.1% saponin and 10% FBS in 1XPBS solution for 1 hour at room temperature. After blocking, the primary anti-ANKZF1 antibody (1:100 dilution was prepared in the blocking solution) was added to the coverslip and incubated at room temperature for 1 hour, then cells were washed thrice with 1XPBS to remove any nonspecific binding. Lastly, secondary antibody Alexa 640 (1:1000 dilution was prepared in the blocking solution) was added for 1 hour at room temperature followed by washing and mounting the coverslip on the slide for imaging.

### ANKZF1 Knockout cell line Generation and Validation

To target the *ANKZF1* gene, sgRNAs were designed by using an online server https://www.vbc-score.org/. Two separate sets of guide RNA (set 1, 5’- TCAAACAGGGAGATCGACGCAGG-3’ and 5’-GCTTGGCCCGAACCGTATAGCGG-3’ and set 2, 5’-TTTCTGCTCAGCCATGTCGCCGG-3’ and 5’- AGAGTGTGATGGCCCACCTCGGG-3’) were selected based on the least off-target effects and high scores to be effective as sgRNAs. The sgRNAs were commercially synthesized and cloned in pSpCas9 (BB)-2A-Puro (PX459) V2.0 (Addgene #62988) plasmid. The cloned sgRNA were transfected in HeLa cells. 24 hours post-transfection cells were supplemented with Puromycin (1µg/ml) for 48 hours, and selected cells were seeded in 96 well plates for the clonal population. Eventually, the surviving populations were screened by the PCR; genomic DNA was isolated from wild type, and both the knockout cell lines and ANKF1 exon- 1 forward and exon- 8 reverse primers were used for PCR amplification. Following PCR-based screening, PCR products were also sequenced by Sanger sequencing, followed by the sequence alignment with the wild-type sequence. Finally, the absence of ANKZF1 was confirmed for knockout clones by western blot using ANKZF1-specific antibody.

### MTT cell viability assay

WT HeLa cells and ANKZF1 knockout HeLa cells were seeded into a 96-well plate at a density of 10,000 cells/well and were allowed to attach for 24 h. One set of both the KO cells (clone 1 and clone 2) was transfected with WT ANKZF1-GFP plasmid, and one set was transfected with Q246L-ANKZF1-GFP plasmid, and were further incubated for 24 hours. Then, all the wells were treated with 100µg/ml Cycloheximide for 24 hours except the untreated control of all three cell lines. All the control and treatment sets were performed in triplicates. Then cells were washed with PBS and 10µl MTT (3-(4,5-dimethylthiazol-2-yl)-2,5-diphenyltetrazolium bromide, Sigma) was added to each well at a final concentration of 0.5 mg/ml and incubated at 37°C for 3 h, which resulted in formazan crystal formation. The formed formazan crystals were dissolved in DMSO and give purple colour. The measurements were taken at 590 nm in a multimode plate reader (Tecan), and results were analysed by GraphPad prism software.

### Western Blot

Cells were lysed by RIPA buffer, protein amounts were quantified, and resolved in polyacrylamide gel electrophoresis and proteins were transferred to PVDF membrane. Membranes were blocked with 5% skimmed milk or 5% BSA dissolved in TBS-T, followed by the primary antibody and secondary antibody incubations. Lastly, blots were developed by using ECL reagents either in Protein Simple Chemidoc system or using the x-ray films.

To detect the PMD protein, whole cell lysate was supplemented with 8M Urea in Laemmli buffer, and sample was heated at 80°C for 10 minutes before loading on gel and then standard western blotting protocol was followed as described above.

### Multiple Sequence analysis

All the multiple sequence alignments were performed in Clustal Omega software and further confirmed in Tcoffee MSA server where protein sequences of the targeted proteins were submitted and checked for the aligned sequences.

### Statistical analyses

All statistical analyses were performed either by using Microsoft Excel or GraphPad Prism using the dataset of three independent repeats. For each experiment at least 25 cells were taken for analysis and at least three biological replicates were done for each experiments. Initially, all datasets were checked for normal distribution, if the datasets were following the normal distribution, then parametric tests like one-way ANOVA with Bonferroni multiple comparison test was used to compare the change difference. However, for the datasets that did not follow a normal distribution, a non-parametric Kruskal-Wallis test was performed to compare the differences. all the datasets were plotted as scatter bar plots, and all the error bars indicate mean ± SEM. ns represents non-significant differences, *P < 0.05, **P < 0.01, ***P < 0.001 ****P < 0.0001.

## Acknowledgments

KM acknowledges the funding support from the Science and Engineering Research Board (SERB), Government of India, for Core Research Grant (SERB/CRG/2022/006517) and SNIoE core funding. MA and Anjali acknowledge the SNIoE PhD fellowship and MA acknowledges the fellowship from ICMR SRF Grant (2021-14421/CMB-BMS). All authors acknowledge the SNU DST-FIST grant [SR/FST/LS-1/2017/59(c)] for the confocal microscopy facility. We thank Rajan Singh for his help at the confocal microscopy facility at SNIoE.

## Author Contribution

The work was conceived by KM. Cell culture, molecular biology and imaging experiments were done by MA. Anjali did the MTT assay and some imaging and biochemical assays. Data were analyzed by MA and KM. KM supervised the work; MA and KM wrote the manuscript.

