## Supplementary Information for "ANKZF1 helps to eliminate stress-damaged mitochondria by LC3-mediated mitophagy"

Running title:

ANKZF1 interacts with LC3 to clear damaged mitochondria

**Supplementary figure legends**

**Figure S1**

**A.** Images of HeLa cells obtained by confocal microscopy after transfection with ANKZF1-GFP and mCherry-Parkin and followed by treatment with H_2_O_2_, rotenone, sodium-azide, and paraquat at concentrations and conditions as mentioned in the materials and methods. In case of rotenone, sodium-azide, and paraquat treatments, cells show some scattered ANKZF1 and Parkin puncta formation, which partially co-localizes to mitochondrial network. **B.** TMRE-uptake based measurement of mitochondrial membrane potential of HeLa cells after treatment with CCCP, rapamycin, H_2_O_2_, rotenone, paraquat, and sodium-azide. Images show reduced TMRE uptake in comparison to control condition in all the treated conditions except H_2_O_2_ treatment. **C.** TMRE uptake of panel **(B)** was calculated by measuring the TMRE intensity and plotted as bar plots. CCCP showed the lowest TMRE uptake while H_2_O_2_ did not show any significant difference from the control cells and the rest of the treatments also showed reduced TMRE uptake. Values represent means ± SEM of N=3, data were not following normal distribution, Kruskal Wallis a non-parametric test with Donn’s multiple comparison test was performed to determine the mean differences, **P < 0.01, ****P < 0.0001.

**Figure S2**

**A.** Western blot showing the presence of Mito-HA-GFP band at ~35kDa in the first lane and mito A53T α-synuclein-GFP band at ~48 kDa in the second lane, but in the third Mito-PMD-GFP band was not detected with anti-GFP antibody, all the samples were prepared in Laemmli buffer. **B.** HeLa cells were transfected with Mito-PMD-GFP, and cells were collected at every three hours intervals of transfection till 21 hours; cells were lysed in 2x Laemmli buffer supplemented with 8M Urea, blot shows the presence of GFP tagged Mito-PMD protein at ~64 kDa. **C.** Schematic workflow of mitochondrial mean branch length measurement using ImageJ.

**Figure S3**

**A.** Left panel in all four rows shows the expression of BFP tagged control and stressor proteins (Mito-BFP, Mito**-**A53T-α-synuclein, and Mito-PMD). Second panel from left, shows the TMRE intensity, cells showing TMRE staining contain healthy mitochondria with intact membrane potential. CCCP-treated cells were used as a negative control for TMRE staining. Expression of Mito-PMD shows significant loss of mitochondrial membrane potential. However, Mito-A53T-α-synuclein expression does not cause much change in the mitochondrial membrane potential. **B.** TMRE uptake shown in panel **(A)** was calculated by measuring the TMRE intensity and plotted. CCCP-treated and PMD-expressing cells show lowest TMRE intensity, Mito**-**A53T-α-synuclein expression does not cause any significant change in TMRE intensity. Values represent means ± SEM of N >3, data were not following normal distribution, Kruskal Wallis a non-parametric test with Donn’s multiple comparison test was performed to determine the mean differences, **P < 0.01, ****P < 0.0001**C.** mCherry-Parkin was co-expressed in HeLa cells expressing either control protein (Mito-BFP), or stressor proteins (A53T-α-synuclein-BFP, or PMD-BFP). In the control panel (Mito-BFP), mCherry-Parkin shows diffused cytosolic expression, but in stressed conditions, lower two panels (A53T α-synuclein BFP and PMD BFP) show a high level of mCherry parkin puncta recruitment on mitochondria indicating increased mitophagic flux due to proteotoxic stress. Inset shows zoomed in boxed regions of the merged panel. **D**. Pearson correlation co-efficient profile of co-localization between mCherry-Parkin and mito-BFP shows significant Parkin recruitment on mitochondria during A53T-Synuclein and PMD stress conditions. Values represent means ± SEM of N>3, dataset was following a normal distribution, One-way ANOVA with Bonferroni’s multiple comparison test was performed to determine the mean differences, ****P < 0.0001.

**Figure S4**

**A.** Cells were transfected with Mito PMD BFP, GFP Parkin and RFP-LC3, followed by the ANKZF1 probing by Anti-ANKZF1 antibody, result shows various co-localized events of Parkin, LC3, and ANKZF1 during mitochondrial proteotoxic stress condition. **B.** HeLa cells were co-transfected with LAMP1 mCherry and ANKZF1 GFP along with mitochondrial stressors protein, top panel shows control condition, ANKZF1 remains in cytosol and no co-localization between ANKZF1 and LAMP1 were observed. Middle and lower panel shows A53T Syn and PMD stressor conditions, in both results ANKZF1 formed puncta and shows significant co-localized events with LAMP1-mCherry. **C**. Pearson’s correlation co-efficient profile of co-localization between LAMP1-mCherry and ANKZF1-GFP shows significant co-localization during A53T-Synuclein and PMD-induced proteotoxic stress condition. Values represent means ± SEM of N>3. Dataset was following a normal distribution. One-way ANOVA with Bonferroni’s multiple comparison test was performed to determine the mean differences, ****P < 0.0001.

**Figure S5**

**A.** ANKZF1 truncation mutants (WT-ANKZF1, Δ210-ANKZF1, Δ330-ANKZF1, and Δ370-ANKZF1) were co-expressed with RFP-LC3 in SHSY-5Y cells in the presence of PMD-induced proteotoxic stress. Stressed cells show ANKZF1 puncta formation by all the truncation mutants except Δ370-ANKZF1. **B**. Pearson’s correlation co-efficient profile of co-localization between RFP-LC3 and ANKZF1-GFP shows Δ370-ANKZF1 fails to localize with LC3. Values represent means ± SEM of N>3, dataset was not following a normal distribution, Kruskal Wallis a non-parametric test with Donn’s multiple comparison test was performed to determine the mean differences, ns: Non significance, ****P < 0.0001. **C.** WT ANKZF1 and F333A-L336A mutant of ANKZF1 were co-expressed with mCherry Parkin and Mito-PMD, showing co-localization (line profile) ANKZF1 and Parkin in both the conditions. **D**. Pearson correlation co-efficient profile of co-localization between mCherry-Parkin and WT-ANKZF1-GFP and LIR mutant ANKZF1-GFP. Values represent means ± SEM of N>3, unpaired students t-test was used to analyze the mean difference, **P < 0.01. **E.** ANKZF1 FL and Ub-DsRed were co-expressed in HeLa cells along with PMD and control protein; top panel shows Mito-BFP control where ANKZF1 remains in cytosol. The middle and lower panel shows PMD and bafilomycin A1 treated PMD conditions, showing ANKZF1 puncta formation and co-localization of ANKZF1 puncta with Ub-DsRed puncta, suggesting ANKZF1 and ubiquitin interaction. **F**. Pearson correlation co-efficient profile of co-localization between Ubiquitin-DsRed and ANKZF1-GFP shows significant co-localization during PMD stress condition. Values represent means ± SEM of N>3, dataset was not following a normal distribution, Kruskal Wallis a non-parametric test with Donn’s multiple comparison test was performed to determine the mean differences, ns: Non significance, ****P < 0.0001.

**Figure S6**

**A.** ANKZF1 recruitment and interaction with Parkin during PMD-induced mitochondrial proteotoxic stress were checked after expressing the N-terminal truncation mutants of ANKZF1. ANKZF1-FL shows puncta formation that co-localizes with mCherry-Parkin puncta during proteotoxic stress (panel A, upper row). Δ330-ANKZF1 truncation mutants also show similar behavior and significantly co-localize with Parkin puncta (panel A, second row from the top) like the full-length protein. Δ370-ANKZF1, Δ410-ANKZF1, and Δ450-ANKZF1 show a diffused cytosolic pattern indicating no interaction with Parkin.  **B**. Pearson correlation co-efficient profile of co-localization between mCherry-Parkin and ANKZF1-GFP show significant loss in the interaction between ANKZF1 and Parkin when at least initial 370 amino acids were deleted. Values represent means ± SEM of N>3, dataset was not following a normal distribution, Kruskal Wallis a non-parametric test with Donn’s multiple comparison test was performed to determine the mean differences, ns: Non significance, ****P < 0.0001. **C**. ANKZF1 recruitment and interaction with Ubiquitin during PMD-induced mitochondrial proteotoxic stress were checked after expressing the truncation mutants of ANKZF1. ANKZF1-FL shows puncta formation that co-localize with Ub-DsRed puncta during proteotoxic stress (panel C, upper row). Δ330-ANKZF1 truncation mutants also show similar behavior and significantly co-localize with Parkin puncta (panel C, second rows from the top) like the full-length protein. Δ370-ANKZF1, Δ410-ANKZF1, and Δ450-ANKZF1 shows a diffused cytosolic pattern indicating no interaction with ubiquitin. **D**. Pearson correlation co-efficient profile of co-localization between Ubiquitin-DsRed and ANKZF1-GFP shows significant loss in the interaction between ANKZF1 and ubiquitin in Δ370-ANKZF1 overexpression. Values represent means ± SEM of N>3, dataset was not following a normal distribution, Kruskal Wallis a non-parametric test with Donn’s multiple comparison test was performed to determine the mean differences, ns: Non significance, ****P < 0.0001.

**Figure S7.**

**A**. I-TASSER predicted-modelled structure of 337-450 residues of ANKZF1, showing formation of α-helices. **B**. Schematic diagram of ANKZF1 domain structure showing the presence of putative UBA domain **C**. Chromatogram showing sequencing confirmation of ANKZF1 LIR mutants generated by site-directed mutagenesis. Left panel shows wild-type sequences with blue highlights at the LIR site, the right panel shows mutant sequences, and red highlights show the mutations being incorporated.

**Figure S8**

**A.** *ANKZF1* gene of both the KO clone cells were sequenced and aligned with the wild type ANKZF1 sequence, showing a gap of around 3 kb suggesting successful deletion of part of the genomic DNA. **B.** Agarose gel electrophoresis of PCR products amplified from genomic DNA of WT HeLa cells and and *ANKZF1* KO cells. PCR was done by using ANKZF1 Exon 1 forward and Exon 8 reverse primers. In the WT HeLa cells, a ~4 kb band was observed, while in KO cells ~1 kb band was observed showing deletion of ~3 kb fragment. **C.** Left panel shows healthy mitochondrial control (Mito-BFP), and PINK1-YFP and mCherry Parkin expression was assessed in wild type and both the *ANKZF1* knockout clones, while right side panel shows PINK1 and Parkin expression during mitochondrial proteotoxic stress Mito-PMD condition. In the control panel, no significant change in the PINK1 or Parkin expression in the KO cells, while in the stressed condition PINK1 and Parkin recruitment on the mitochondria was significantly increased in all three conditions, no overall cell to cell variability was observed.

**Figure S9.**

**A.** Bar plot showing the cell viability percentage of wt HeLa and *ANKZF1* KO clones, first three bars showing untreated control, bar 4-6 showing cells exposed with translation inhibitor cycloheximide (100µg/ml) for 24 hours, where *ANKZF1* KO cells showing sensitivity to cycloheximide treatment. Bar 7 and 8 shows *ANKZF1* KO clones were rescued with the overexpression of wild type ANKZF1 and treated with cycloheximide (100µg/ml) for 24 hours, showing overexpression of WT ANKZF1 helps in the cell survival. Last 2 bars represent the conditions where *ANKZF1* KO clones were rescued with the overexpression of tRNA hydrolase deficient mutant Q246L ANKZF1 and treated with cycloheximide (100µg/ml) for 24 hours, here the cell death was not rescued by overexpression of mutant ANKZF1. Values represent means ± SEM of N =3, data was following a normal distribution, one-way ANOVA with Bonferroni correction for multiple comparison was performed to determine the mean differences, ns: non-significant differences, ***P < 0.001, ****P < 0.0001. **B.** Q246L ANKZF1 was overexpressed and checked its interaction with Parkin, during mitochondrial proteotoxic stress Mito-PMD in wt HeLa and *ANKZF1* Knockout clones, in every condition Q246L ANKZF1 GFP was able to co-localize with mCherry Parkin. **C.** Pearson’s correlation co-efficient profile of Q246L ANKZF1 GFP with mCherry Parkin in wt HeLa and both the *ANKZF1* KO clones showing no significant changes. **D.** Q246L ANKZF1 was overexpressed and checked its interaction with RFP-LC3, during mitochondrial proteotoxic stress Mito-PMD in wt HeLa and *ANKZF1* Knockout clones, in every condition Q246L ANKZF1 GFP was able to co-localize with RFP-LC3. **E.** Pearson’s correlation co-efficient profile of Q246L ANKZF1 GFP with RFP-LC3 in wt HeLa and both the *ANKZF1* KO clones showing no significant changes. Values represent means ± SEM of N >3, data were not following normal distribution, Kruskal Wallis a non-parametric test with Donn’s multiple comparison test was performed to determine the mean differences, ns: non-significant differences.

**Figure S10**

**A.** The mitophagy rate of WT HeLa cells and *ANKZF1* KO cells were compared by using pH dependent Mt-Keima protein as explained in the results. KO clone 2 cells showed functional mitophagy events but relatively lowered mitophagy events in comparison to WT HeLa cells when PMD protein was expressed to induce mitochondrial proteotoxic stress. **B.** Keima560/Keima488 Ratiometric analysis suggests PMD expressed WT-Hela cells showing significantly higher 560/488 ratio in comparison to the KO cells corresponding high mitophagy rate. **C.** ANKZF1 was overexpressed in ANKZF1 KO cells and Mt-Keima based mitophagy turnover was analysed, in PMD expressed cells, Keima 561 signal behaving like WT HeLa cells, suggesting reversal of phenotype in presence of ANKZF1. Values represent means ± SEM of N >3, data were not following normal distribution, Kruskal Wallis a non-parametric test with Donn’s multiple comparison test was performed to determine the mean differences, *P < 0.05, **P < 0.01. **D.** WT HeLa cells and ANKZF1 KO cells were transfected with tf-LC3 (GFP and RFP tagged LC3) along with control and PMD-BFP overexpression in mitochondria. In both the cell lines (WT and KO) the overall autophagy flux is not compromised in presence of PMD proteotoxic stress, almost same number of red LC3 puncta were observed suggesting functional autophagy.

**Figure S11.**

**A and B.** Mt-Keima was expressed in WT HeLa cells and ANKZF1 KO cells (clone 1 and clone 2) followed by the CCCP treatment to induce mitophagy, followed by the confocal microscopy. Results show in WT cells Keima488 to Keima560 shift is higher than the KO cells, suggesting higher mitophagy rate in WT cells in comparison to KO cells. Values represent means ± SEM of N >3, data were not following normal distribution, Kruskal Wallis a non-parametric test with Donn’s multiple comparison test was performed to determine the mean differences, *P < 0.05, **P < 0.01.

**Supplementary table 1.**

**List of plasmid constructs used in this work**

| **Construct Name** | **Vector Backbone** | **Source** | **Addgene Number** |
| --- | --- | --- | --- |
| EBFP2-Mito-7 | EBFP2 | Addgene | 55248 |
| mCherry-Parkin | mCherry-C1 | Addgene | 23956 |
| Mito HA-GFP | pcDNA3.1 | Addgene | 67486 |
| mCherry-Mito-7 | mCherry | Addgene | 55102 |
| ptf-LC3 | EGFP-C1 | Addgene | 21074 |
| mt-mKeima | pHAGE | Addgene | 131626 |
| RFP-LC3 |  | Received as a gift from Professor Oishee Chakrabarti’s lab, SINP, Kolkata |  |
| Mito-A53T-Synuclein-GFP | EGFPN1 | This study |  |
| Mito-PMD-GFP | EGFPN1 | This study |  |
| Mito-A53T-Synuclein-BFP | EBFP | This study |  |
| Mito-PMD-BFP | EBFP | This study |  |
| ANKZF1-FL-GFP | pcDNA3.1 | This study |  |
| Δ210-ANKZF1-GFP | EGFPN1 | This study |  |
| Δ330-ANKZF1-GFP | EGFPN1 | This study |  |
| Δ370-ANKZF1-GFP | EGFPN1 | This study |  |
| ANKZF1-333AXXA336-GFP | EGFPN1 | This study |  |
| ANKZF1-366AXXA369-GFP | EGFPN1 | This study |  |
| ANKZF1-495AXXA498-GFP | EGFPN1 | This study |  |
| Q246L ANKZF1-GFP | EGFPN1 | This study |  |
| Ub G76V GFP | EGFPN1 | Addgene | 11941 |
| Ub G76V DsRed | DsRedN1 | This Study |  |
| LAMP1-emiRFP670 | emiRFP670 | Addgene | 136570 |
| LAMP1-mCherry | mCherry | This Study |  |

**Supplementary table 2.**

**List of reagents and antibody used in this paper.**

| **Reagents and Antibody** | **Made** | **Cat number** |
| --- | --- | --- |
| Lipofectamine^TM^ 2000 | Thermo | 11668027 |
| H_2_O_2_ | MP Bio | Addgene |
| Rotenone | Sigma-Aldrich | R8875 |
| Rapamycin | Sigma-Aldrich | 553210 |
| CCCP | Sigma-Aldrich | C2759 |
| Paraquat | Sigma-Aldrich | 856177 |
| Na-Azide | Himedia | MB075 |
| Cycloheximide | MP Bio | 194527 |
| Thiazolyl blue tetrazolium bromide (MTT) | Sigma | M2128 |
| MitoTracker Far Red | Invitrogen | M22426 |
| TMRE | Invitrogen | T669 |
| Anti-ANKZF1 | Sigma | HPA035208 |
| Anti-GFP | Generated in Lab |  |
| Anti-LC3 | Cell signalling technology | 3868T |
| Anti-Parkin | Cell signalling technology | 2132S |
| Anti-Actin | Elabscience | E-AB-40517 |
| Alexa-647 | Invitrogen | A21245 |

**Supplementary table 3**

| **Primer Name** | **Primer Sequence** |
| --- | --- |
| ANKZF1 EcoRI Forward | 5’CCCGAATTCATGTCGCCGGCTCCAGATGCAGCCCCGGCTCC3’ |
| 211 ANKZF1 EcoRI Forward | 5’ CCCGAATTCATGGCTGCAGCTGGGCACTTTGCTGGTG 3’ |
| 330-ANKZF1 XhoI Forward | 5’ GTTCTCGAGATGCCCACCTTCCAAGAGCTAC 3’ |
| 370-ANKZF1 XhoI Forward | 5’ GTTCTCGAGATGGAGGAGAGAAAGAAGCCTAC 3’ |
| 410-ANKZF1 XhoI Forward | 5’ GTTCTCGAGATGTTTAGGTAGAGTTGGAGC 3’ |
| ANKZF1 No stop KpnI Reverse | 5’ CCCGGTACCCGGGAAGAGGGCCTCCCTGCCTGACGGCGATG 3’ |
| ANKZF1 F333A and L336A SDM Forward | 5’ AGACCCACCGCCCAAGAGGCCCAGCGTGTG 3’ |
| ANKZF1 F333A and L336A SDM Reverse | 5’ CACACGCTGGGCCTCTTGGGCGGTGGGTCT 3’ |
| ANKZF1 W366A and V369A SDM Forward | 5’ CAGACACACGCCAAAACAGCCAGAGAGGAG 3’ |
| ANKZF1 W366A and V369A SDM Reverse | 5’ CTCCTCTCTGGCTGTTTTGGCGTGTGTCTG 3’ |
| ANKZF1 W495A and L498A SDM Forward | 5’ CCAGAGCTCGCCAATGCAGCCCTTGCTGCT 3’ |
| ANKZF1 W495A and L498A SDM Reverse | 5’ AGCAGCAAGGGCTGCATTGGCGAGCTCTGG 3’ |
| ANKZF1 Q246L Forward | 5’ GGGCACAGCCCTGGGGCTTCGGG 3’ |
| ANKZF1 Q246L Reverse | 5’ CCCGAAGCCCCAGGGCTGTGCCC 3’ |
| ANKZF1 Exon 1 Forward | 5’ CGCTGCTCCGTAGTGACGG 3’ |
| ANKZF1 Exon 8 Reverse | 5’ CCCGGTACCCGGATATCCCAAAGTCGGGGATCC 3’ |
| ANKZF1 sgRNA 1 Forward Oligo | 5’ CACCGCTTGGCCCGAACCGTATAG 3’ |
| ANKZF1 sgRNA 1 Reverse Oligo | 5’ AAACCTATACGGTTCGGGCCAAGC 3’ |
| ANKZF1 sgRNA 2 Forward Oligo | 5’ CACCGAGTGTGATGGCCCACCTC 3’ |
| ANKZF1 sgRNA 2 Reverse Oligo | 5’ AAACGAGGTGGGCCATCACACTC 3’ |
| ANKZF1 sgRNA 3 Forward Oligo | 5’ CACCGTTTCTGCTCAGCCATGTCGC 3’ |
| ANKZF1 sgRNA 3 Reverse Oligo | 5’ AAACGCGACATGGCTGAGCAGAAAC 3’ |
| ANKZF1 sgRNA 4 Forward Oligo | 5’ CACCGTCAAACAGGGAGATCGACGC 3’ |
| ANKZF1 sgRNA 4 Reverse Oligo | 5’ AAACGCGTCGATCTCCCTGTTTGAC 3’ |
