## Supplementary figures and images for "ANKZF1 helps to eliminate stress-damaged mitochondria by LC3-mediated mitophagy"

### Supplementary Figure 1

#### Figure S1

A

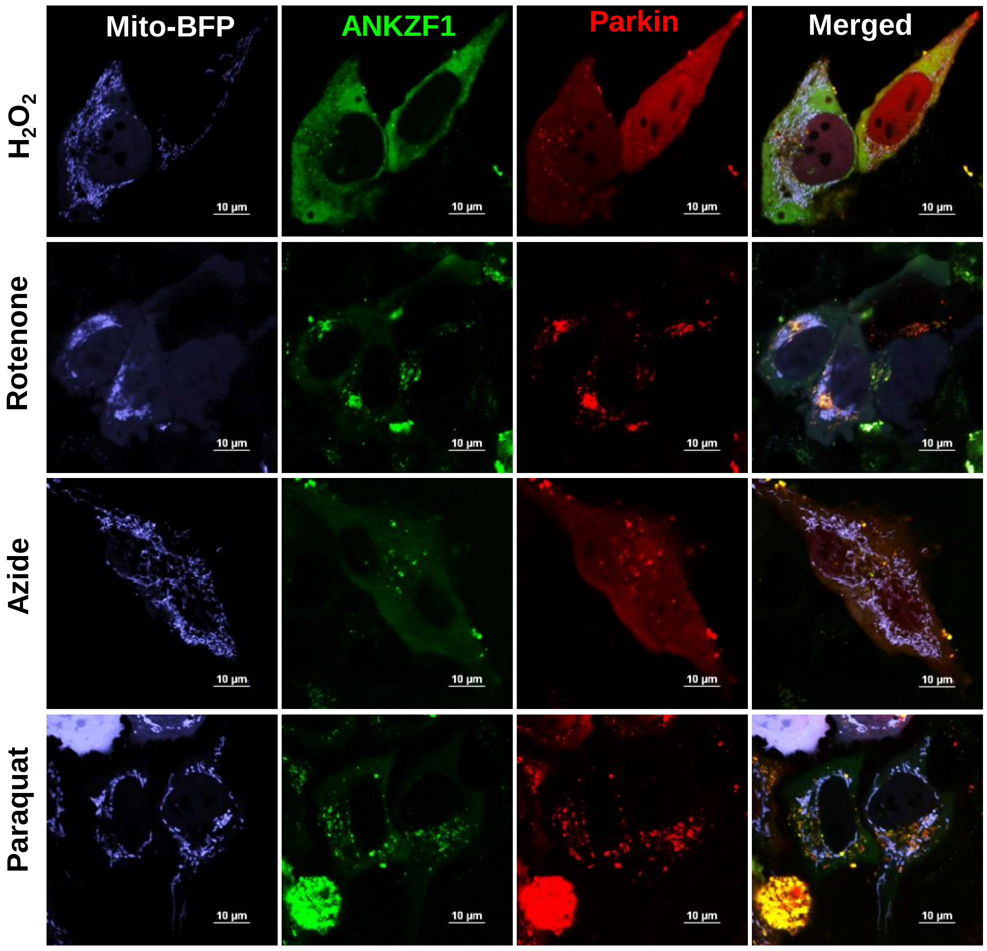


B

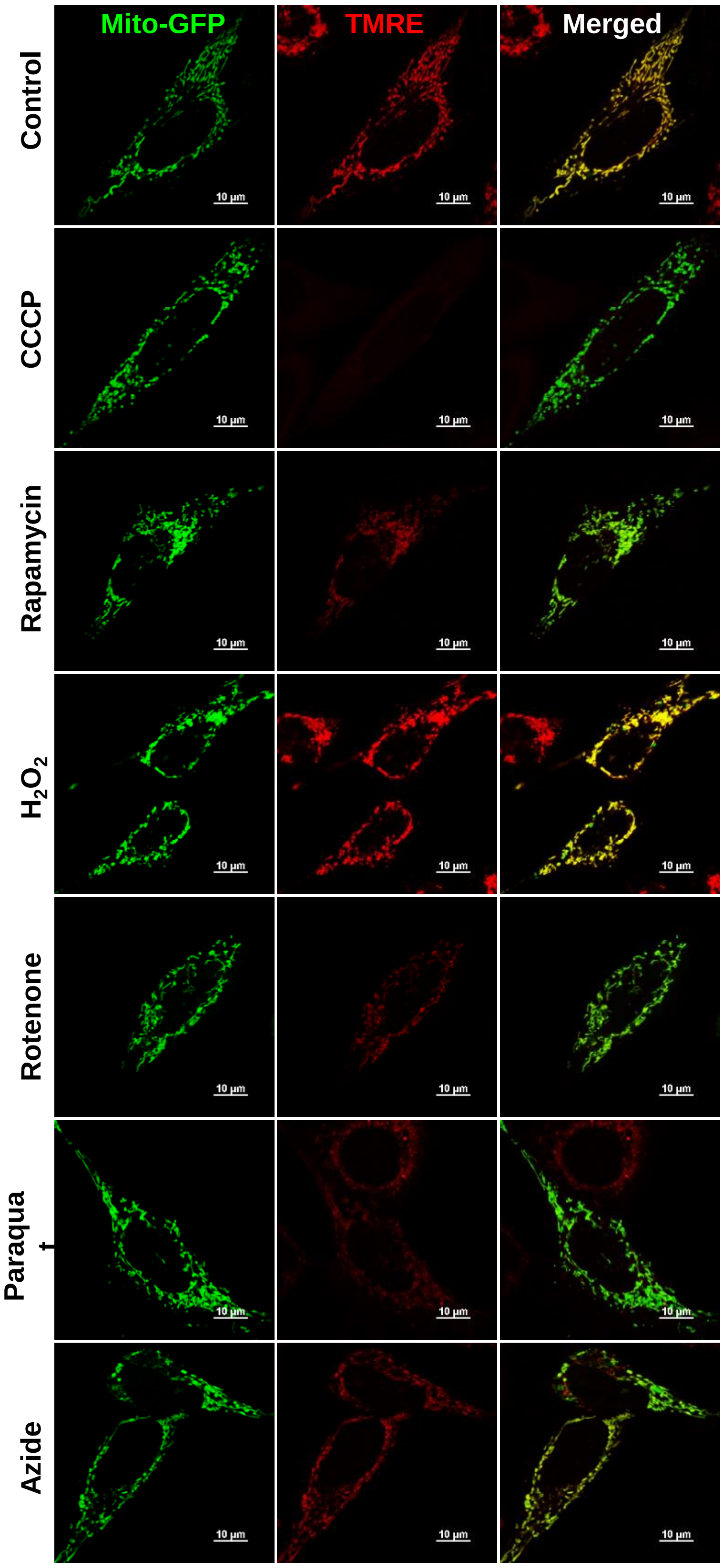


C

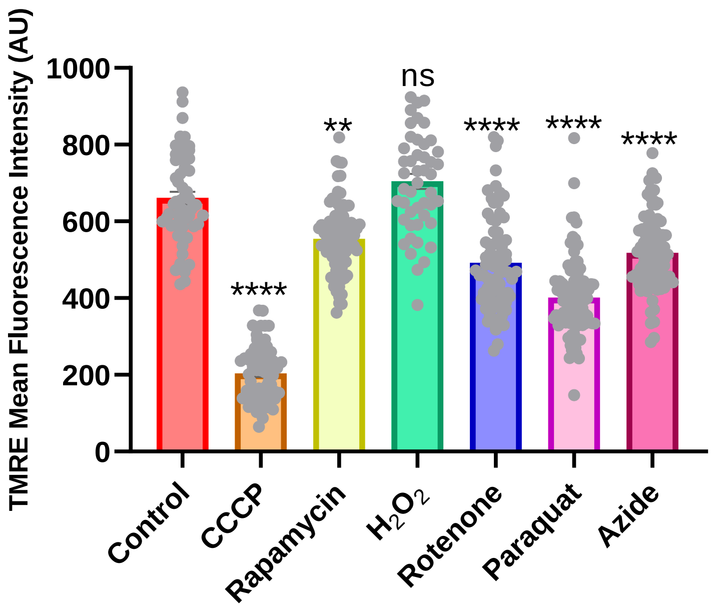

### Supplementary Figure 2

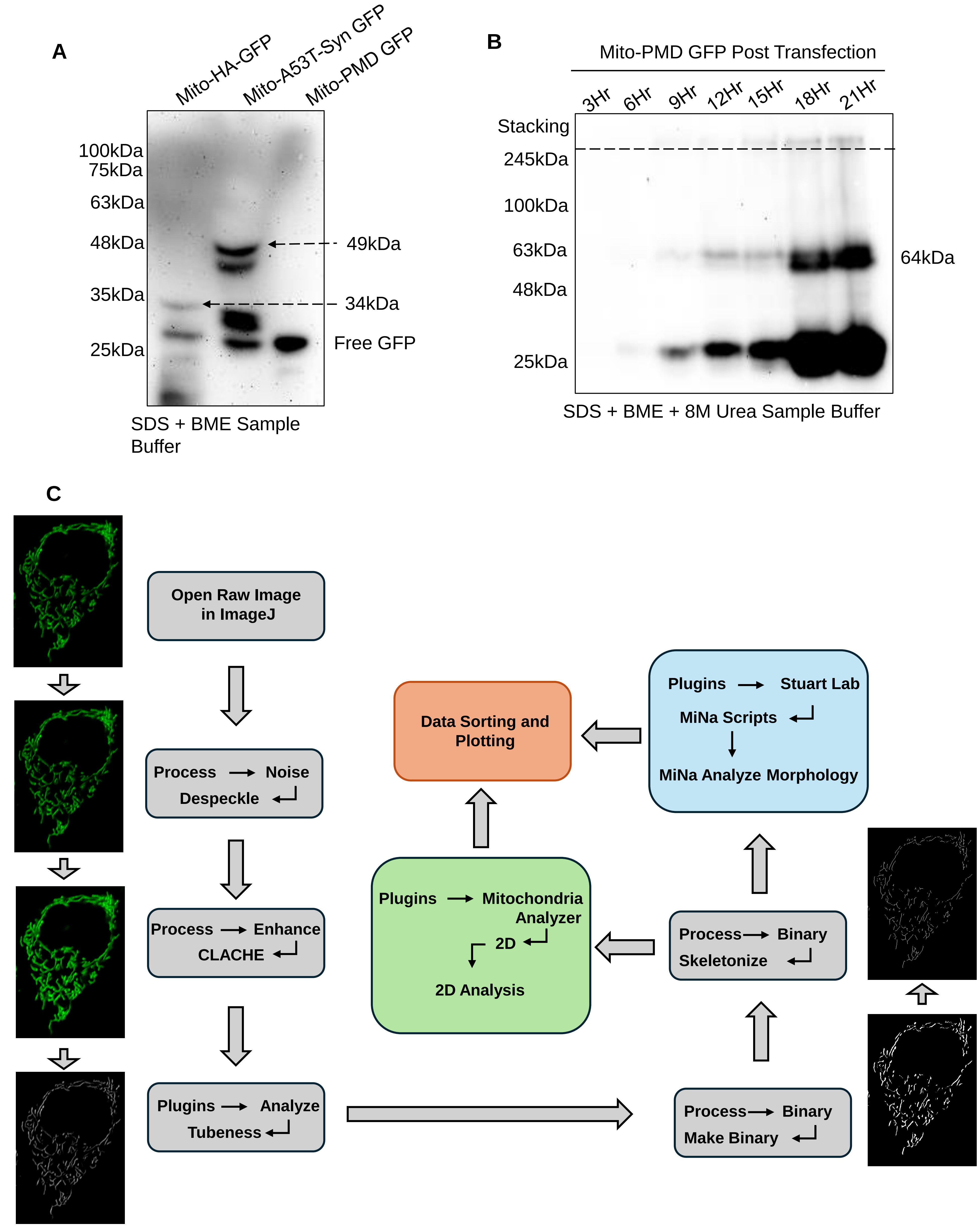

### Supplementary Figure 3

##### Figure S3

A

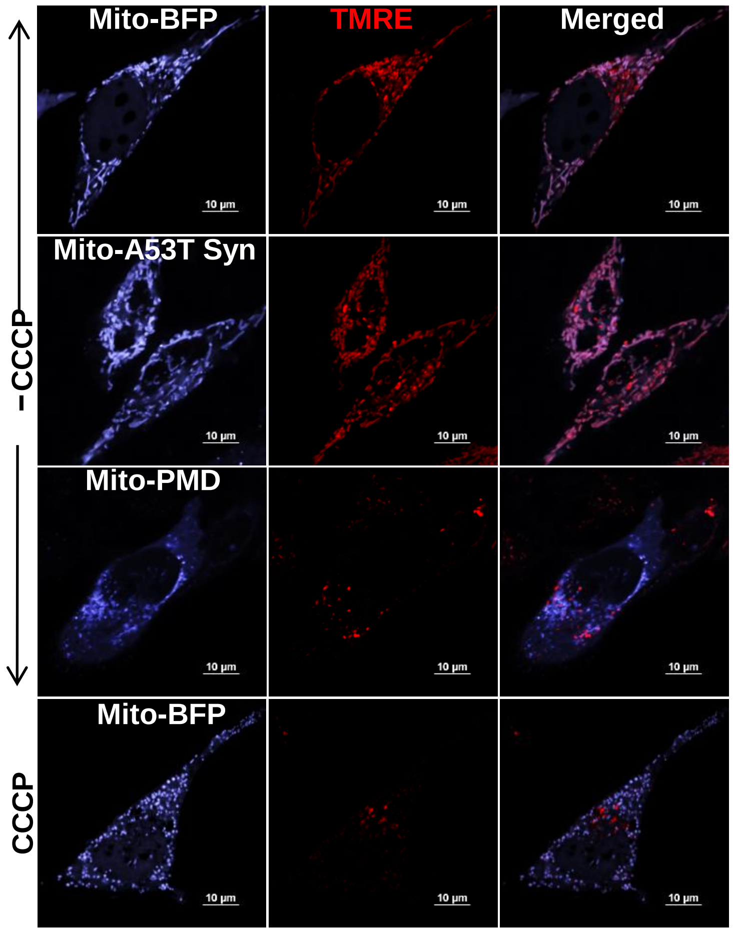


B

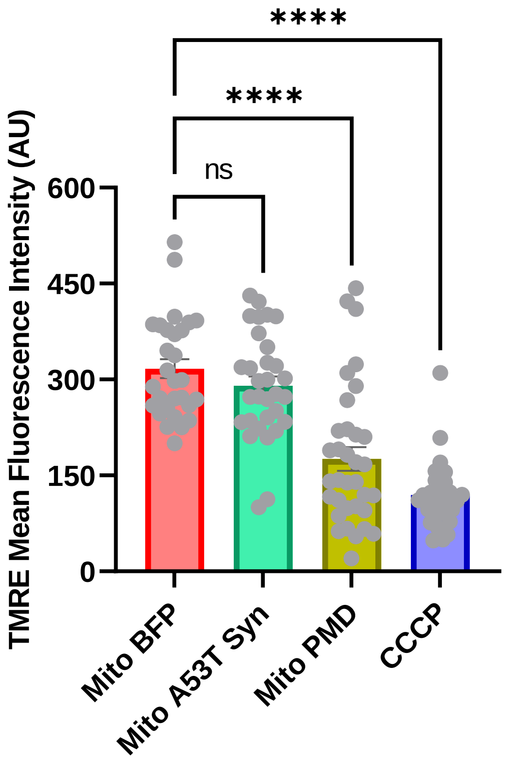


C

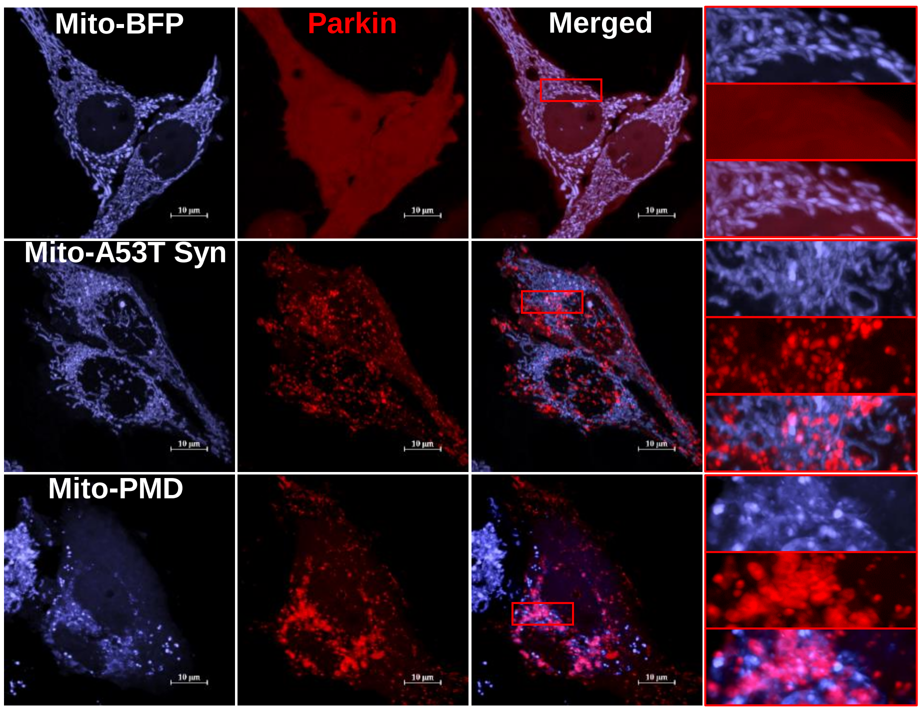


D

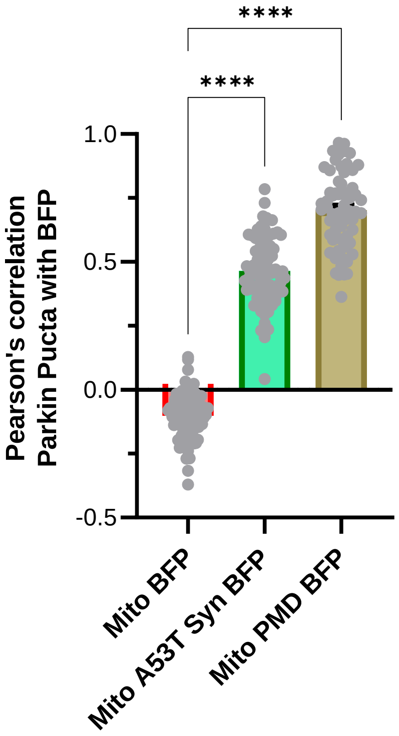

### Supplementary Figure 4

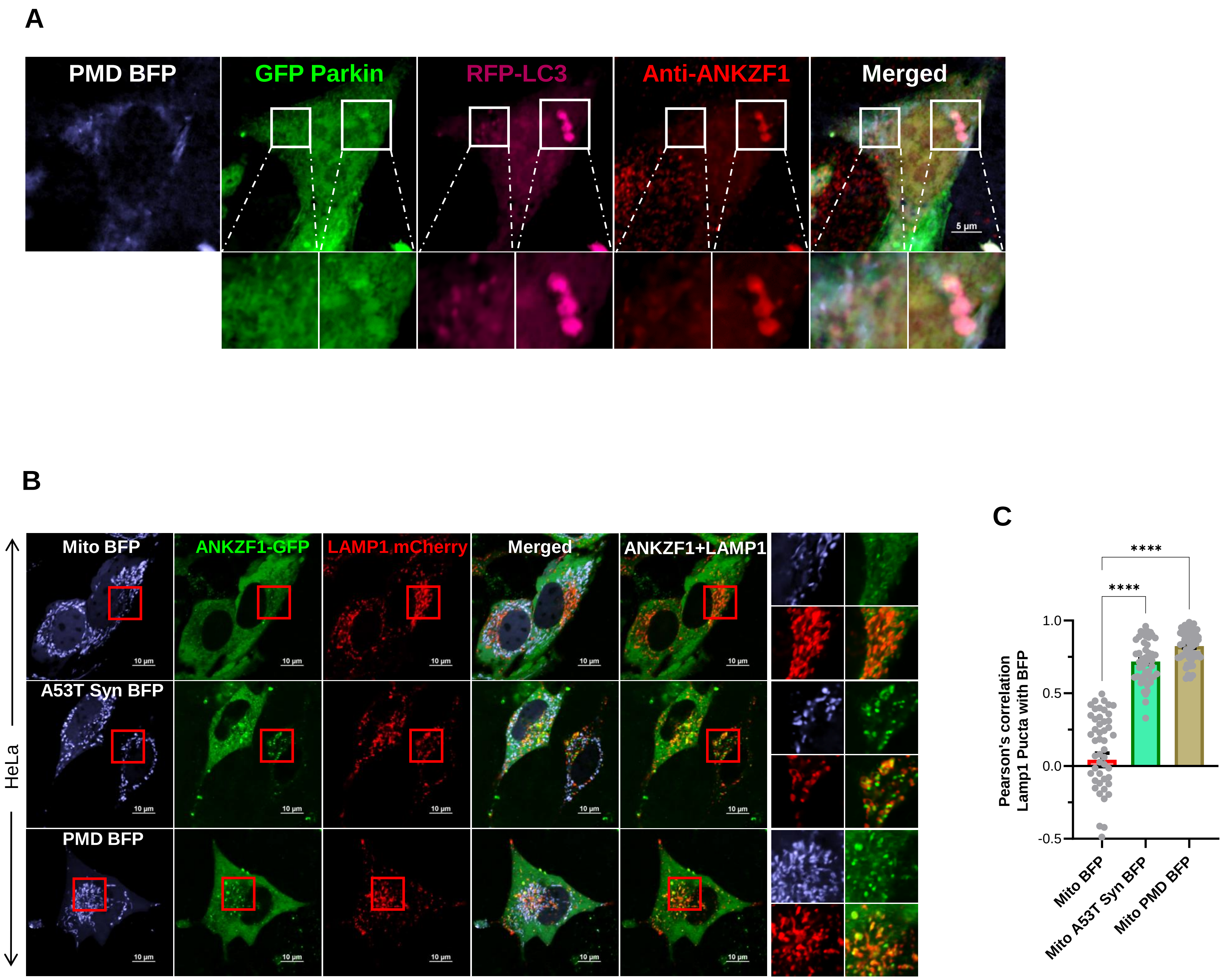

### Supplementary Figure 5

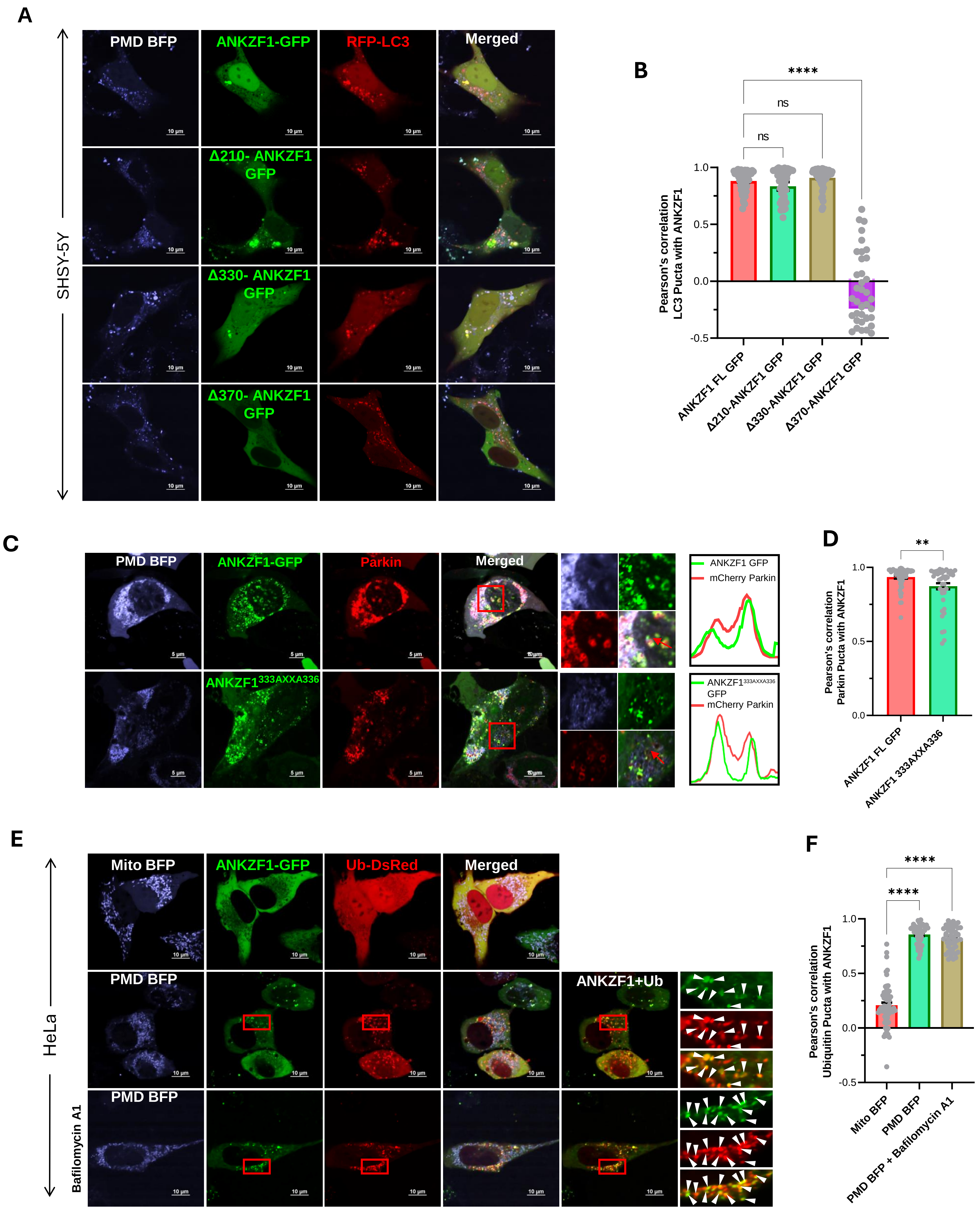

### Supplementary Figure 6

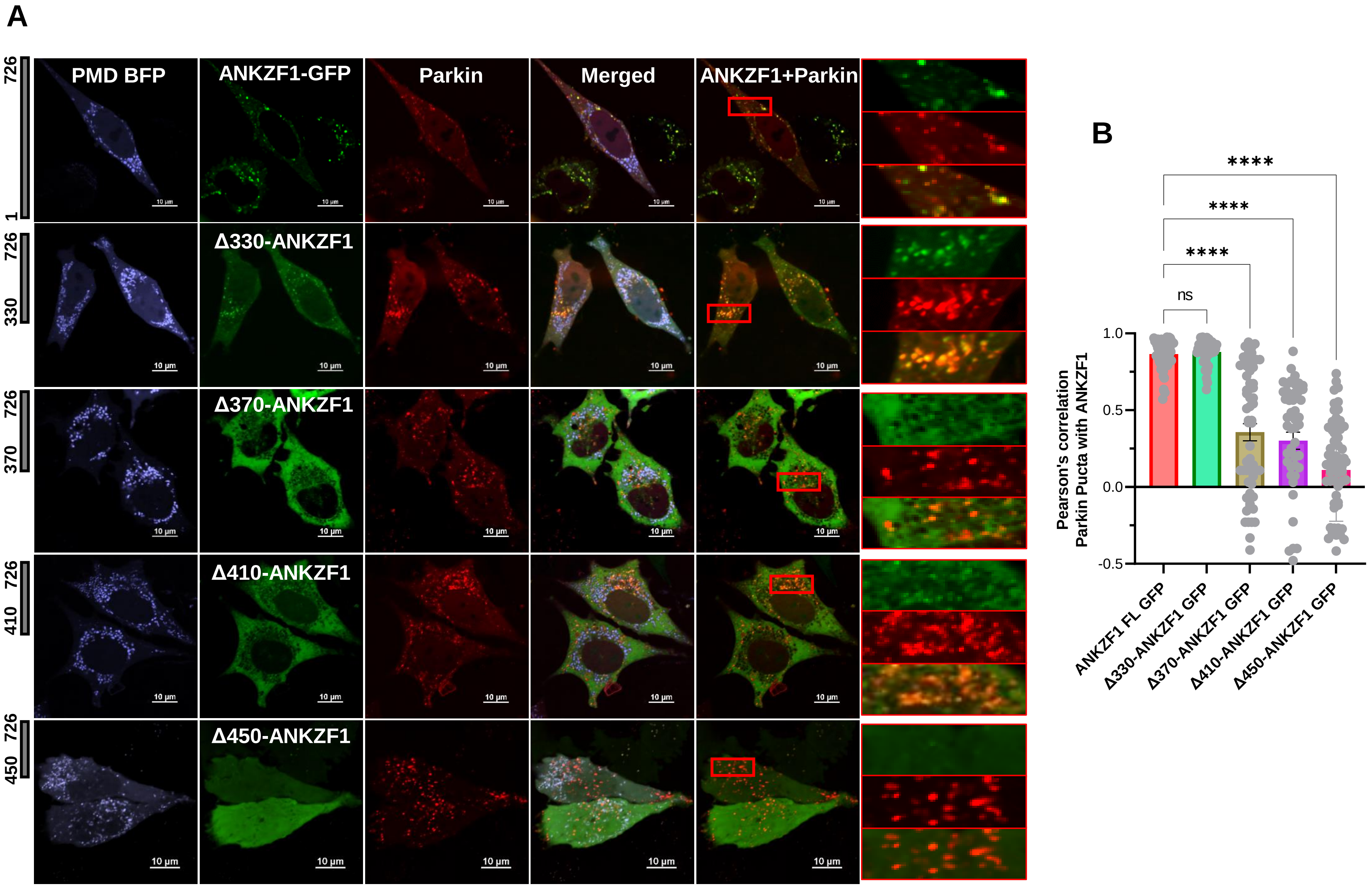

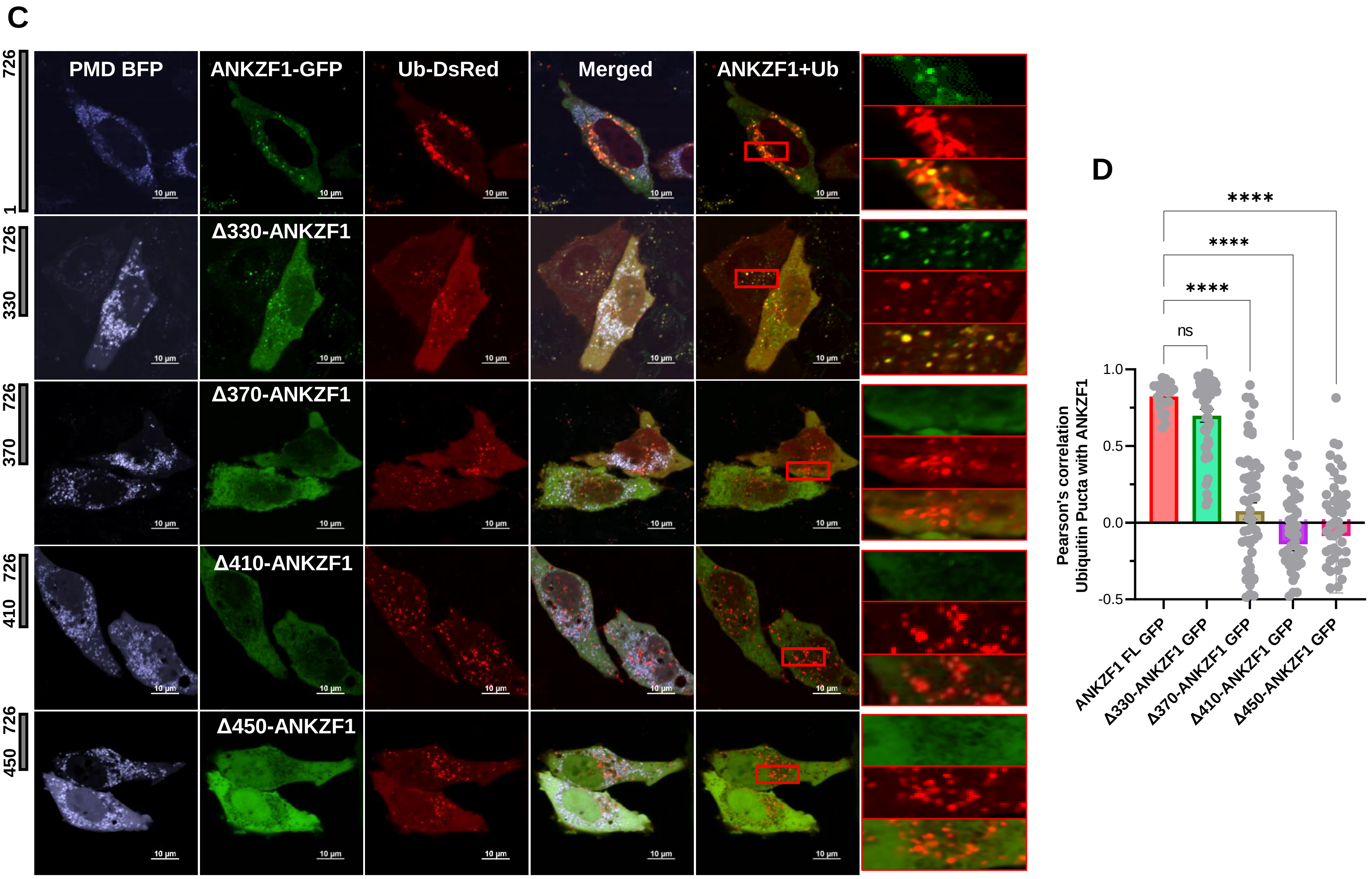

### Supplementary Figure 7

#### Figure S7

A

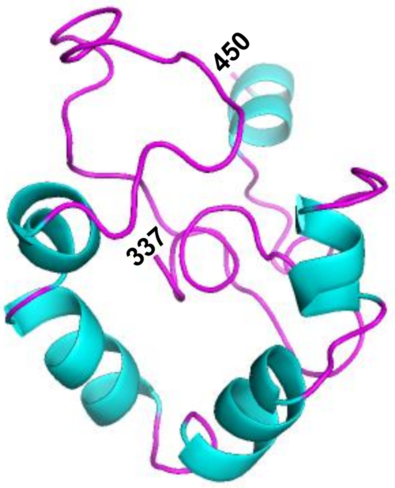


B

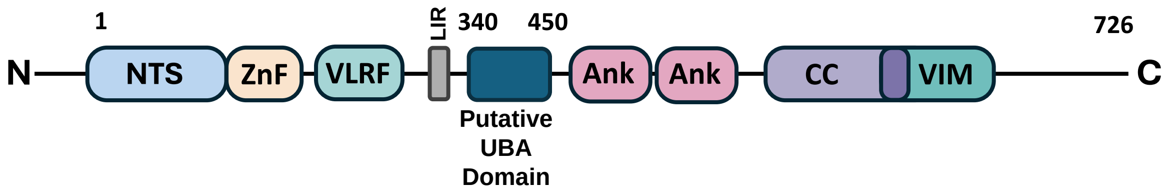


C

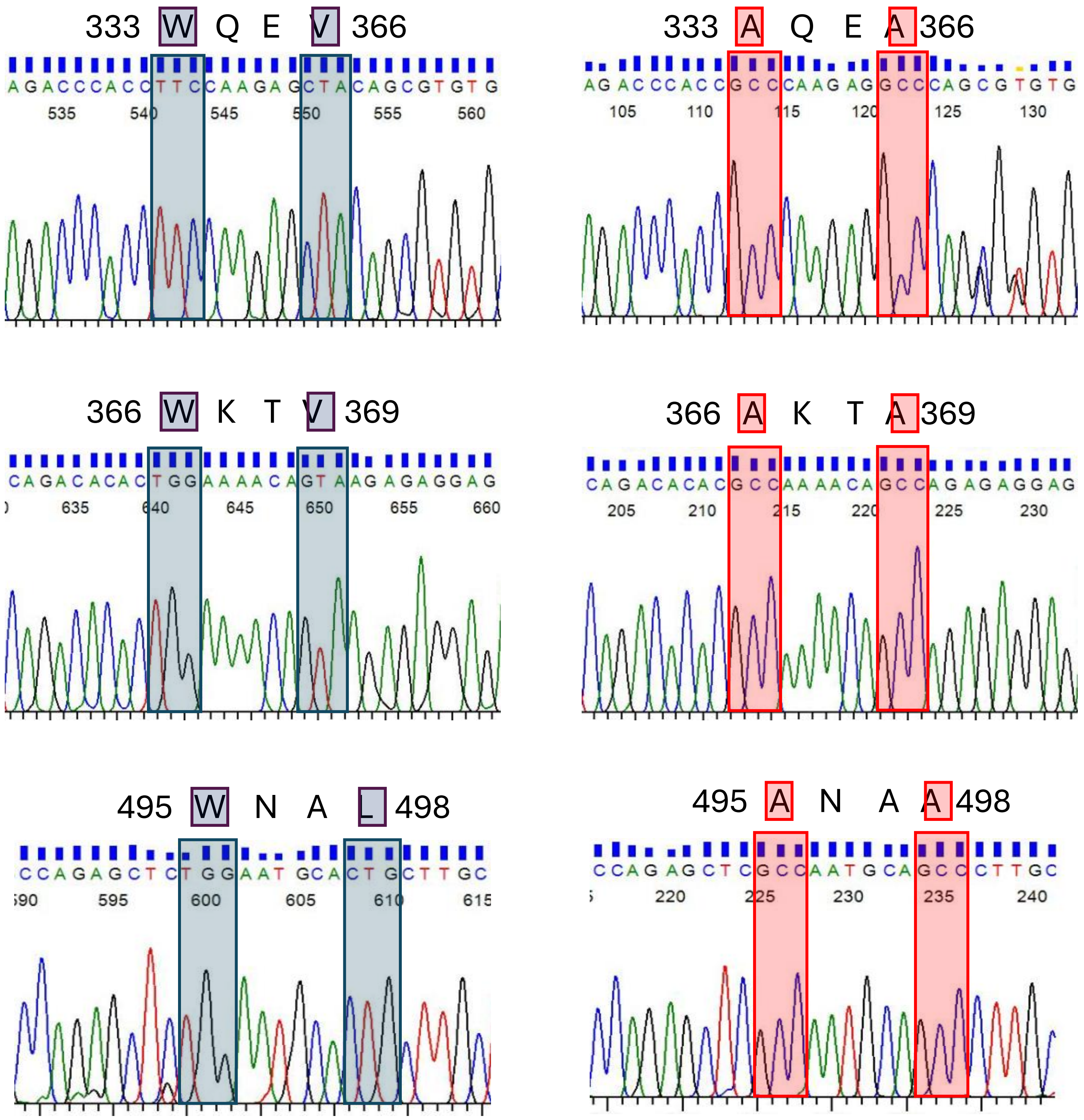

### Supplementary Figure 8

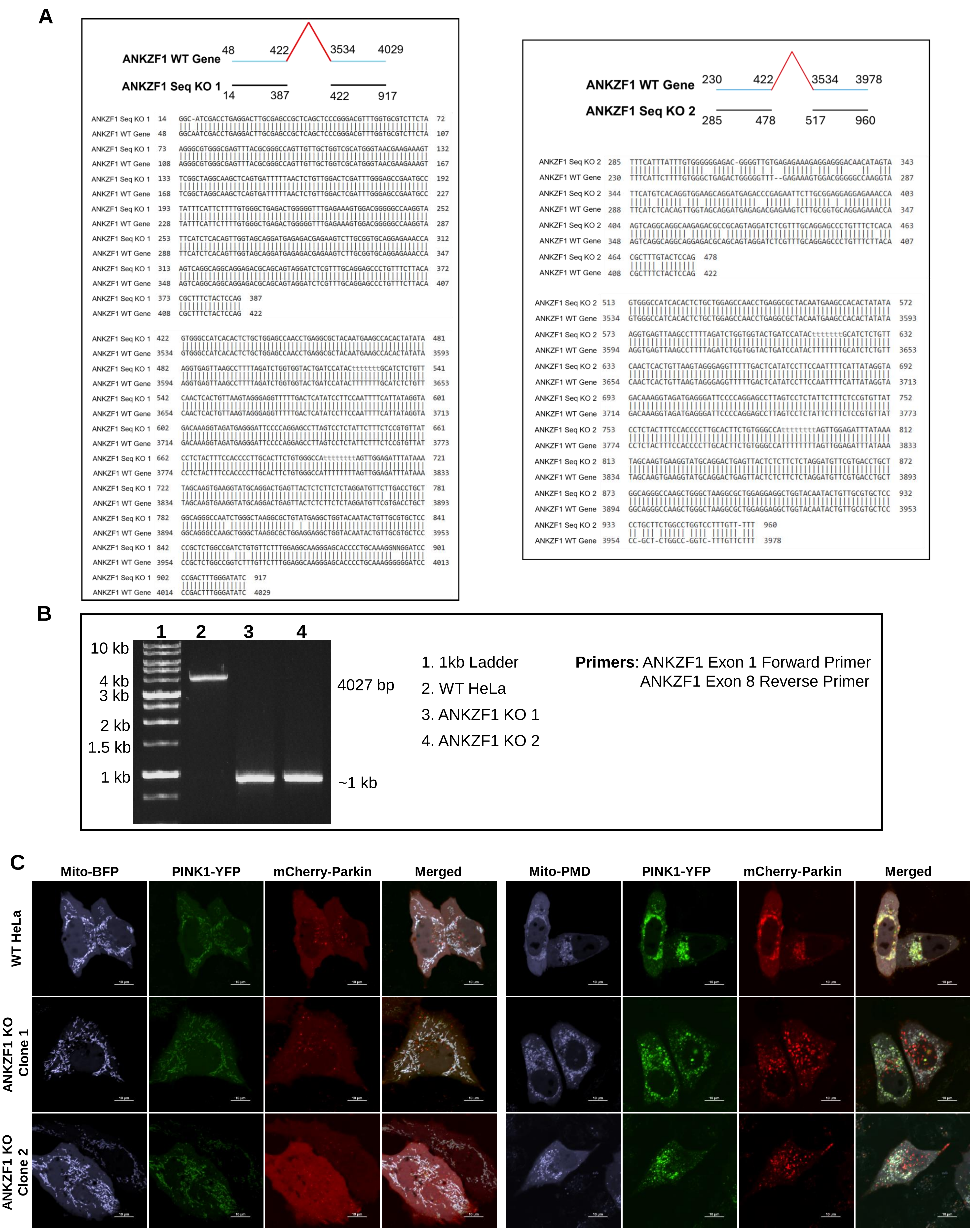

### Supplementary Figure 9

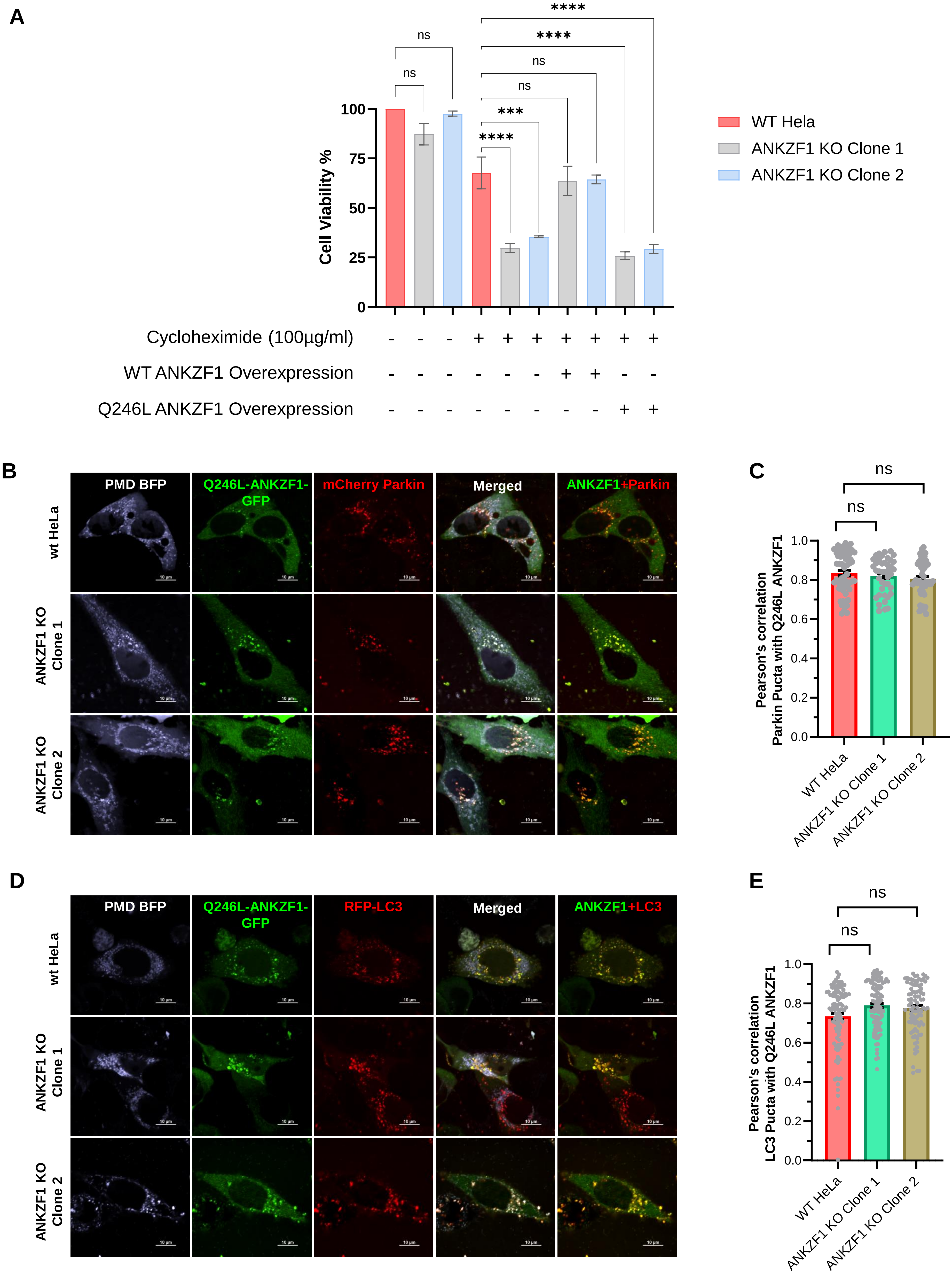

### Supplementary Figure 10

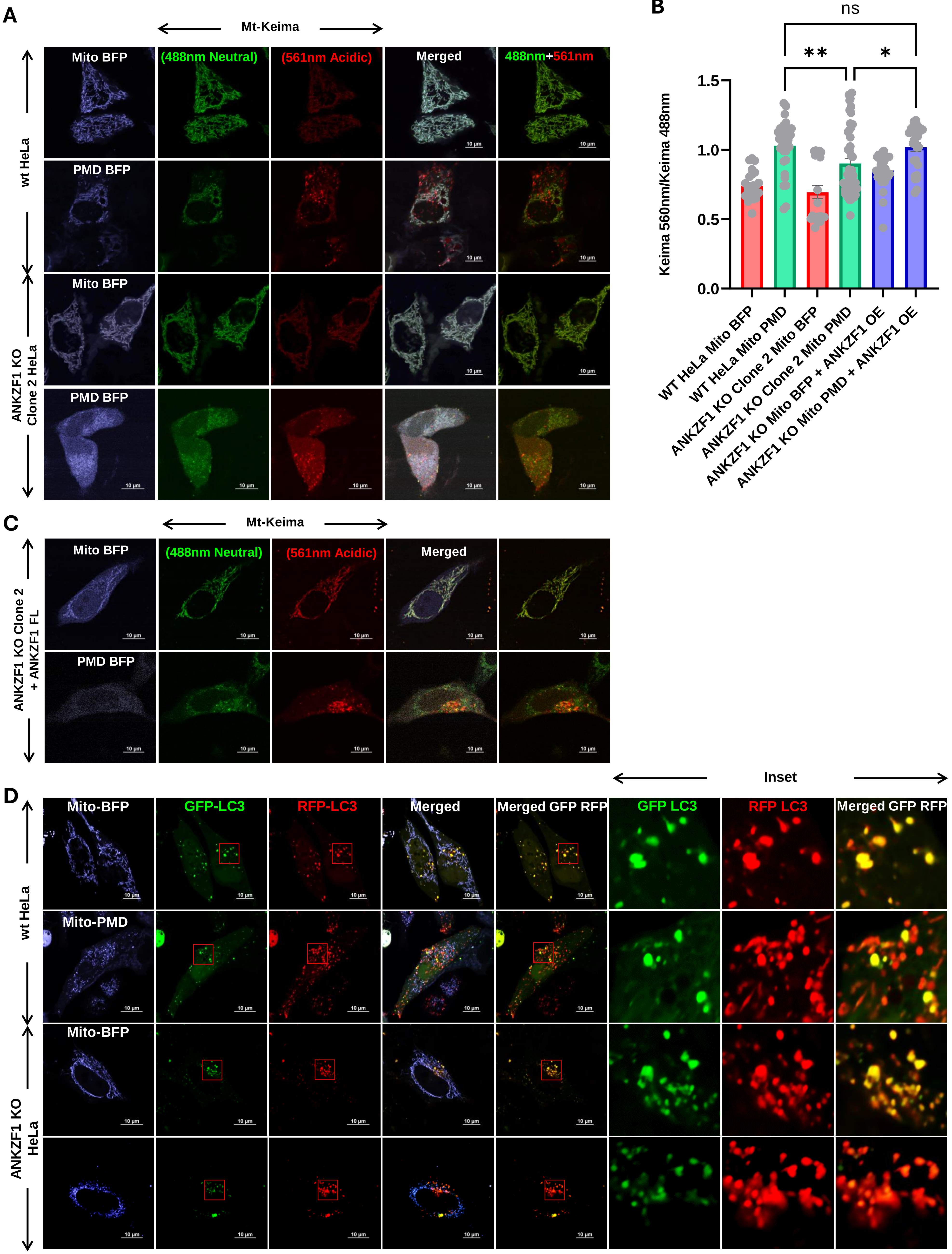

### Supplementary Figure 11

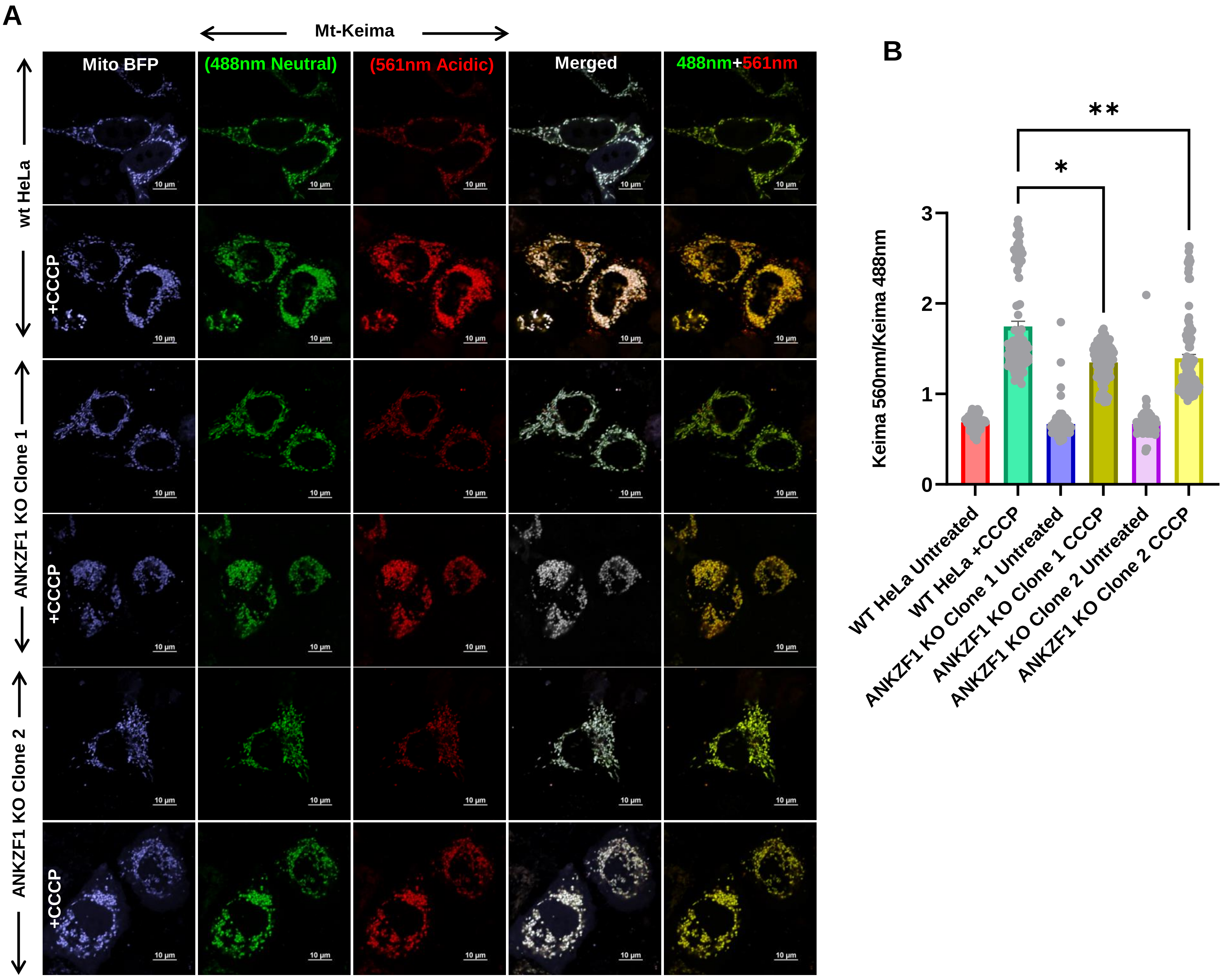

### Supplementary Figure 12

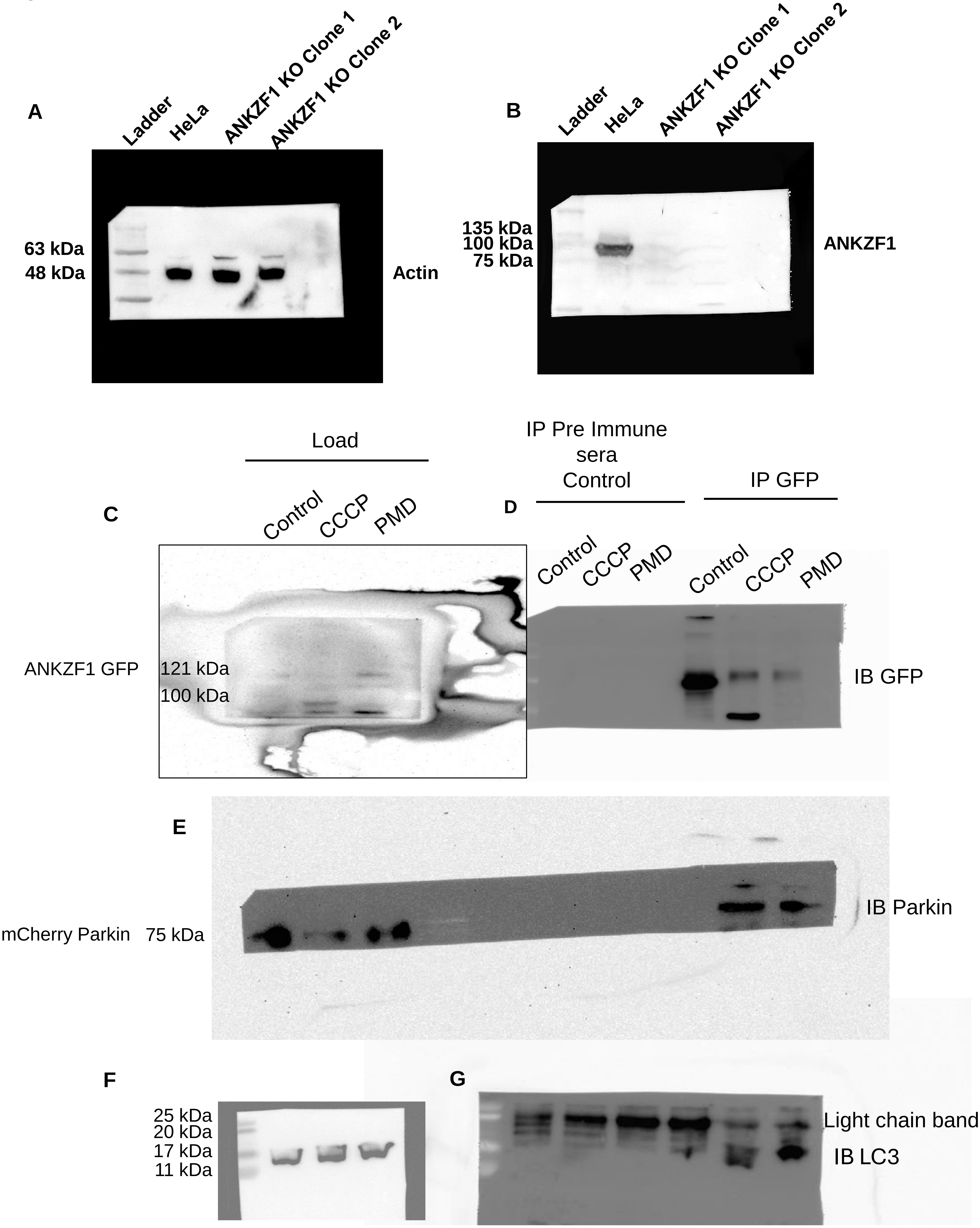
